# Tissue- and sex-specific DNA damage tracks aging in rodents and humans

**DOI:** 10.1101/2022.11.28.518087

**Authors:** Axel Guilbaud, Farzan Ghanegolmohammadi, Yijun Wang, Jiapeng Leng, Alexander Kreymerman, Jacqueline Gamboa Varela, Jessica Garbern, Hannah Elwell, Fang Cao, Elisabeth M. Ricci-Blair, Cui Liang, Seetharamsingh Balamkundu, Charles Vidoudez, Michael S. DeMott, Kenneth Bedi, Kenneth B. Margulies, David A. Bennett, Abraham A. Palmer, Amanda Barkley-Levenson, Richard T. Lee, Peter C. Dedon

## Abstract

DNA damage causes genomic instability underlying many human diseases. Traditional approaches to DNA damage analysis provide minimal insights into the spectrum of disease-driving DNA lesions and the mechanisms causing imbalances in damage formation and repair. Here we used untargeted mass spectrometry-based adductomics^1^ to discover 114 putative DNA lesions and modifications consistently detected in humans and two independent analyses in rats, showing species-, tissue-, age-, and sex-biases. As evidence of methodologic rigor, 10 selected adductomic signals were structurally validated as epigenetic marks: 5-MdC, 5-HMdC, 5-FdC; DNA damage products: *N^2^*-CMdG, 1,*N*^6^-*ε*dA, 3,*N*^4^-*ε*dC, M^1^dG, *O*^6/^*N*^2^- MdG, and 8-Oxo-dG; and established analytical artifacts: cyclobutane dimers of 2’- deoxycytosine. With steady-state levels of putative DNA adducts integrating multiple cell types in each tissue, there was strong age-dependent variation for many putative adducts, including *N^2^*-CMdG, 5-HMdC, and 8-Oxo-dG in rats and 1,*N*^6^-εdA in human heart, as well as sex biases for 67 putative adducts in rat tissues. These results demonstrate the potential of untargeted adductomic analysis for defining DNA adducts as disease determinants, assigning substrates to DNA repair pathways, discovering new metabolically-driven DNA lesions, and quantifying inter-individual variation in DNA damage and repair across populations.

## Main text

### Refinement and validation of untargeted adductomics

Traditional targeted approaches to analyzing DNA adducts and structural alterations have revealed the existence of dozens of types of damage caused by both endogenous metabolic processes and exogenous environmental agents^2,3^. However, targeted approaches have yielded little information about the true spectrum of DNA adducts and lesions driving aging and disease in human tissues. Here we took an untargeted low-resolution mass spectrometry-based “adductomic” approach^1, 4–8^ to characterize known DNA lesions and modifications and discover new ones unique to individual rat and human tissues and the behavior of these damage products as a function of age and sex. The adductomics approach involves an untargeted discovery phase in which the 2’-deoxynucleoside components of hydrolyzed DNA are analyzed by multiple reaction monitoring (MRM) of collision-induced dissociation (CID) products losing 2’- deoxyribose (*m/z* 116) in 1 Dalton increments across the range of *m/z* 225-525 (*i.e.,* “stepped MRM”). This approach provides greater sensitivity (≤ 10 fmol; *vide infra*) than simple neutral loss scanning. After filtering data to remove known artefacts and other noise (**Fig. 1a**), the remaining putative DNA adduct signals serve as a basis set for subsequent MRM-based LC-MS/MS analysis of individual DNA samples to quantify the putative adducts. For all studies in rat, mouse, and human tissues, care was taken to avoid adventitious damage during DNA isolation and processing, with addition of antioxidants and deaminase inhibitors^9^.

**Figure 1.**
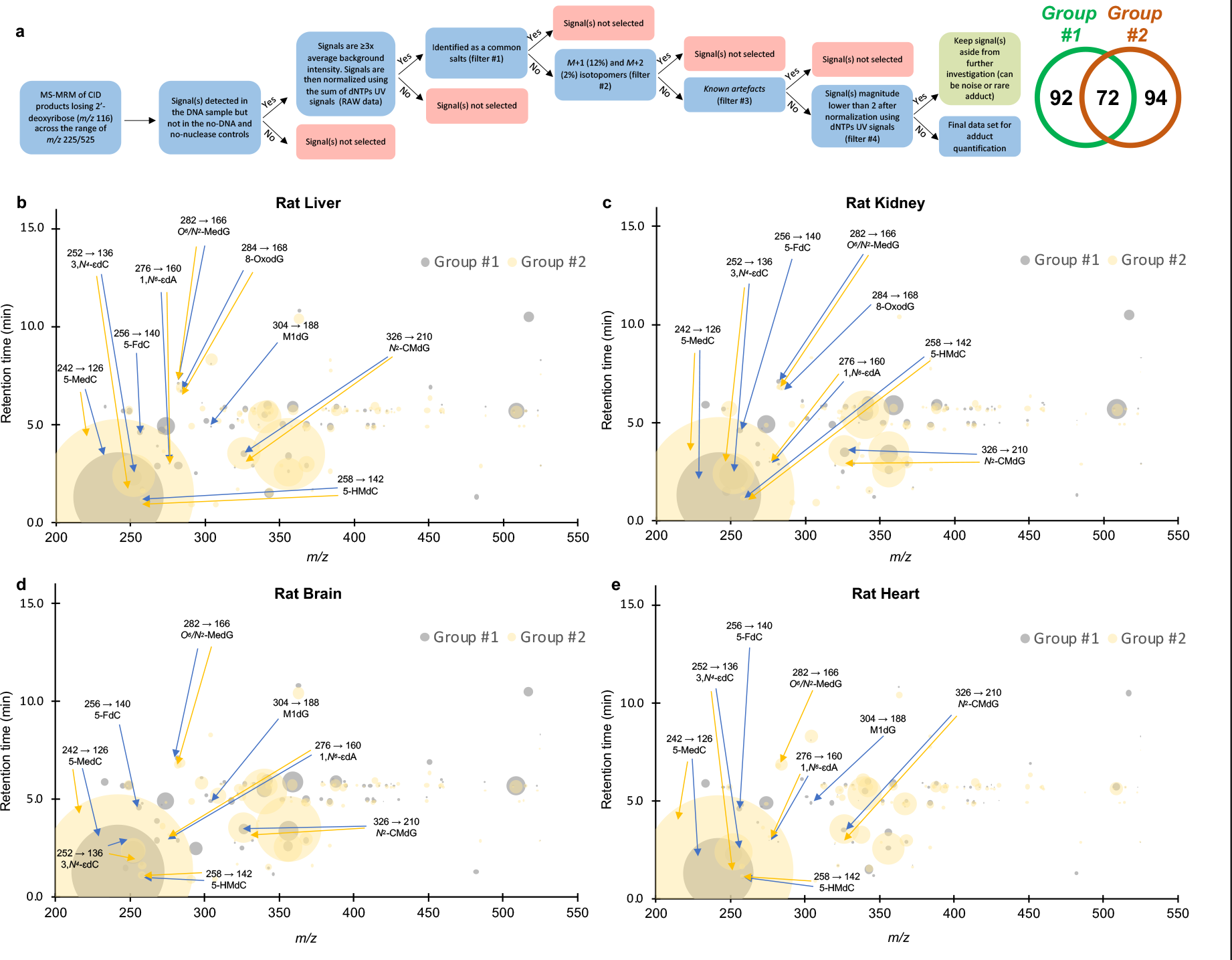
Adductomic analysis in two different sets of rat tissues. Adductomics involves two stages of data acquisition: (1) discovery of putative adducts by stepped MRM analysis of pooled DNA samples and (2) subsequent targeted analysis of a high-confidence set of adductomic signals representing putative DNA adducts in each tissue sample. (**a**) For the first step of identifying putative DNA adducts, the stepped MRM signals must be curated to remove artifacts, noise, and other spurious signals. Here we developed a decision tree using different analytical filters to remove common salts, isotopomers, known artifacts, and potential noise (signal intensity <2) to obtain a final data set of DNA adducts for subsequent targeted analysis. The Venn diagram shows the final sets of 92 and 94 putative adducts identified in discovery studies for tissues from two different rat cohorts, yielding a total set of 114 putative DNA adducts and modifications for subsequent quantitative analysis in each sample. (**b**) Using the curated set of adducts, DNA adductome profiles were generated for individual DNA samples from rat liver (**b**), kidney (**c**), brain (**d**), and heart (**e**) in rats ranging in age from 1 to 26 months for rat group #1 (blue) and 4 to 26 months for rat group #2 (yellow) months. DNA adducts that were further identified using chemical standards are labelled with names and CID transitions. Each DNA adduct, defined or putative, is represented by its retention time (min, y-axis), mass-to-charge ratio (*m/z*, x-axis) and relative abundance (size of the bubble). Signals from the four canonical DNA nucleobases are not represented in the plot. Blue bubbles and arrows are DNA adduct signals observed in group #1 while yellow are those observed in group #2.

**Supplementary Table 1** details the adductomics discovery phase using DNA isolated from four tissues from groups of four male and four female Brown-Norway rats ranging age from 1- to 26-months-old, in two completely independent studies performed on different mass spectrometers with different rat cohorts. To provide the broadest coverage of adduct variation, DNA samples from each tissue were pooled across all ages to create four samples subjected to adductomics discovery analysis. The 300 mass spectrometer signals from adduct discovery for each organ were reduced to 92 and 94 mass spectrometer signals in the two independent rat studies and assigned as putative DNA adducts and modifications on the basis of several filtering criteria (**Fig. 1a**): (1) MS fragmentation released a mass consistent with 2’- deoxyribose, (2) signals were detected in at least one tissue at a level that was at least three- times the average background signal intensity and were detected in all individual samples of each rat or human tissue, (3) signals were not detected in no-DNA and no-nuclease controls (*i.e.,* all reagents except DNA or nuclease/phosphatase), (4) signals representing salts (*e.g.,* Na^+^, K^+^) were removed, (5) signals representing putative isotopomers of stronger signals (*m/z* +1 and +2) were removed, (6) known artifacts were removed, (7) signals with normalized signal intensities ≤ 2.0 and not shared between the two independent analyses were removed. It is important to note that the mass spectrometer signals could represent co-eluting mixtures of chemicals with the same *m/z* value and cannot be assumed to represent individual 2’- deoxyribonucleosides until the signals are structurally validated. The power of the adductomics method lies in the ability to discover all detectable damage products and modifications in cells or tissues and then quantify their variance among tissues and conditions, with subsequent structural analysis confirming the identity of the putative DNA damage products or modifications as the first step in mechanistic understanding of disease-driving genetic toxicology.

As evidence of the rigor of the discovery method, 72 of putative adducts (63%) were shared between the two independent cohorts of rats analyzed with two different mass spectrometer systems (Agilent 6490 and 6495) (**Fig. 1a**, **Supplementary Table 1**). We then attempted to assign chemical structures to each of the 114 putative adducts according to literature data^2, 5– 7, 10–15^, subsequent high-resolution mass spectrometric analysis of HPLC fractions, or by comparison to synthetic standards. The adductomic analysis revealed 66 previously undescribed putative 2’-deoxyribonucleoside species as well as 48 species with fragmentation and *m/z* values consistent with known damage products (average normalized signal intensities and tentative chemical identities in **Supplementary Table 1** and **Extended Data Figure 1**). These results are summarized in the bubble plots in **Figure 1**, depicting the identity and quantity of putative DNA damage products in liver (**Fig. 1a**), kidney (**Fig. 1b**), brain (**Fig. 1c**), and heart (**Fig. 1d**) tissues from the two rat cohorts (blue and yellow bubbles).

That the untargeted adductomic method revealed true DNA damage products and modifications is supported by confirmation of the structures of 10 signals as known DNA lesions based on comparison to chemical standards (**Extended Data Fig. 1c**). For example, we detected a putative adduct with *m/z* of 242→126 in all tissues, which was validated as the epigenetic marker 5-methyl-2’-deoxycytidine (5-MdC; **Fig. 2**). The fact that the signal intensity for the *m/z* of 242→126 5-MdC modification is several orders-of-magnitude higher than other putative adducts in all tissues (**Fig. 1, Supplementary Table 1**) is consistent with its presence as ∼1% of all dC in the human genome^16, 17^. Also consistent with the quantitative rigor of the method, we detected an age-dependent signal with *m/z* 326→210 at much lower levels than 5-MdC and identified it as *N*^2^-carboxymethyl-2’-deoxyguanosine (*N*^2^-CMdG) (**Fig. 2**, **Supplementary Table 1**) – an advanced glycation end-product resulting from the reaction between glyoxal and 2’-deoxyguanosine – by chemical synthesis of a standard for comparison by high-resolution LC-MS analysis (**Extended Data Fig. 1c**). Similar structural validation was achieved for DNA damage arising from oxidation (8-hydroxy-2’- deoxyguanosine, 8-Oxo-dG) and alkylation (1,*N*^6^-etheno-2’-deoxyadenosine, 1,*N*^6^-*ε*dA; 3,*N*^4^- etheno-2’-deoxycytosine, 3,*N*^4^-*ε*dC; *N*^1^-methyl-2’-deoxyguanosine, M^1^dG; *O*^6^- or *N*^2^-methyl- 2’- deoxyguanosine, *O*^6^-or *N*^2^-MdG), as well as physiological epigenetic marks (5- hydroxymethyl-2’-deoxycytidine, 5-HMdC; 5-formyl-2’-deoxycytidine, 5-FdC). The rigor of the method was further validated with a putative adduct having an *m/z* 455→339 (**Supplementary Table 1**). High-resolution MS^4^ Orbitrap mass spectrometric analysis revealed the structure of this adduct as the cyclobutane dimer of 2’-deoxycytosine (dC=dC; **Extended Data Fig. 2**). That this adduct co-eluted with dC is consistent with its formation as an artifact of ionization and not as a pre-existing damage product in DNA.

**Figure 2.**
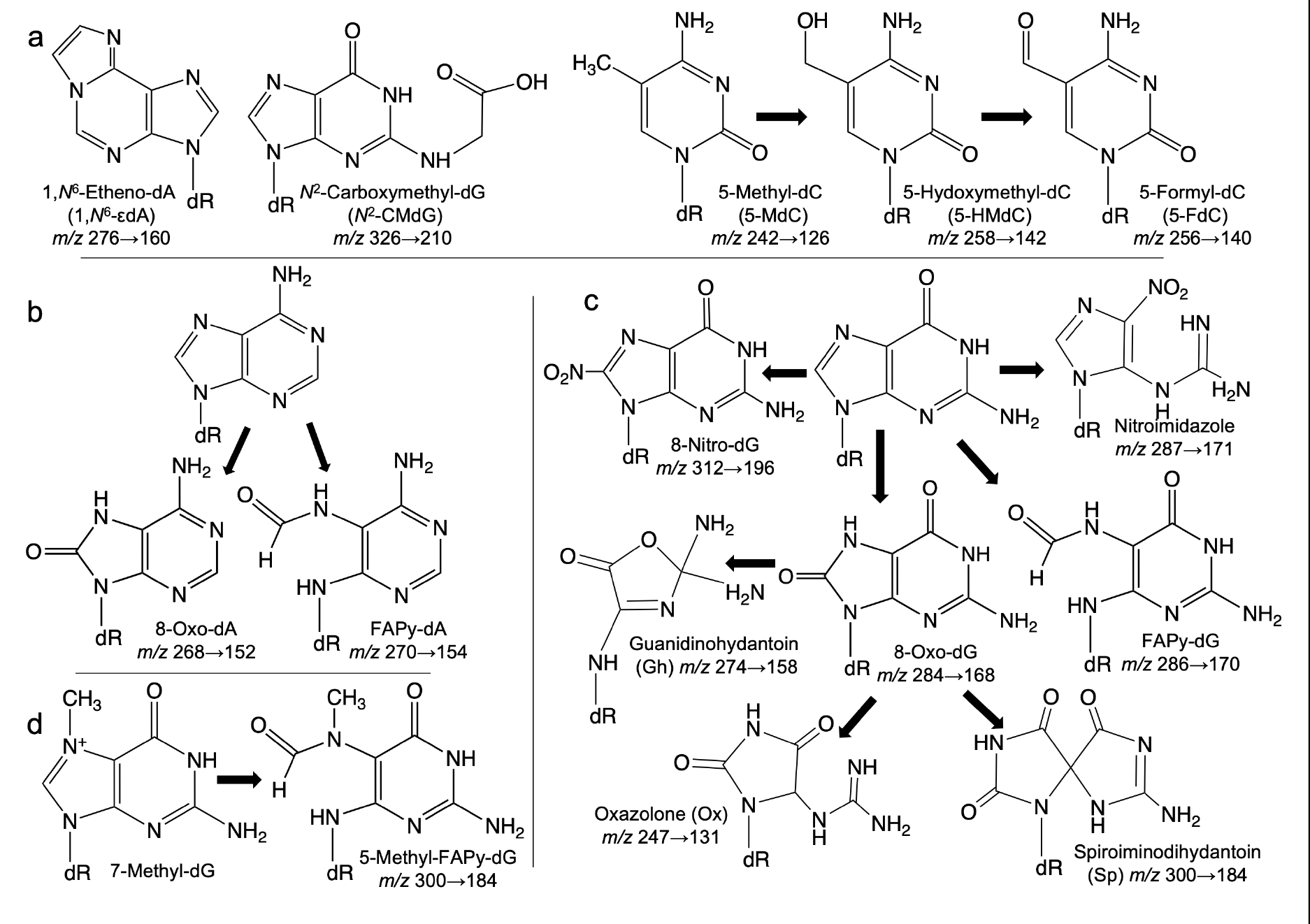
Known DNA damage products and modifications found or predicted among adductomic signals. (**a**) Using synthetic standards, adductomic signals were validated as the DNA damage products *1,N^6^*-*ε*dA, *N^2^*-CMdG, and 8-Oxo-dG (panel **c**), among others, as well as the epigenetic marks 5-MdC and 5-HM-dC. (**b-d**) Oxidation of canonical 2’- deoxynucleosides and DNA damage products leads to a host of secondary products predicted to occur *in vivo,* including FAPy-dA (*m/z* 271.1→155.1), guanidinohydantoin *(m/z* 274.1→158.1), 8-nitro-dG (*m/z* 313.1→197.1), and oxazolone (*m/z* 247.1→131.1). See **Supplementary Table 1** for adductomic signal data.

Either due to lack of formation or levels below detection limits, we did not detect DNA halogenation products (*e.g*., 5-chloro-2’-deoxycytidine, 8-chloro-2’-deoxyguanosine, 8- chloro-2’deoxyadenosine, 2-chloro-2’-deoxyadenosine) or DNA deamination products 2’- deoxyxanthosine (dX; from 2’-deoxyguanosine), 2’-deoxyinosine (dI; from 2’- deoxyadenosine), and 2’-deoxyuridine (dU; from 2’-deoxycytidine). dX is relatively unstable and depurinates during mass spectrometric ionization^9^, while pyrimidine nucleosides such as dU do not ionize well^18^, which may explain our inability to detect these damage products in the set of 114 detected species. Rough upper boundaries for the limits of quantification (LOQ) by the stepped MRM adductomics method can be inferred from targeted analyses of 8-Oxo-dG, *N^2^*-CMdG, 5-HMdC and 1,*N*^6^-*ε*dA by isotope-dilution LC-MS (**Extended Data Fig. 1**), which revealed LOQ values as follows (compound, fmol injected, lesions per 10^9^ nt): 8-Oxo-dG, 0.9, 49; *N^2^*-CMdG, 3.8, 209; 5-HMdC, 4.2, 231; 1,*N*^6^-*ε*dA, 2.6, 143. The 114 species identified by this validated adductomic method are thus very likely to be DNA damage products and modifications meriting further characterization given their strong associations with tissue, sex, and age in mice, rats, and humans.

### Tissue- and sex-specific DNA damage and modification in rats

The observation of 114 putative DNA adducts identified in two independent studies of four tissues from male and female rats that were 1- to 26-months-old raised the question of tissue-, sex-, and age-specificity of the individual adduct species. At the simplest level, a comparison of adductomic signal intensities (**Extended Data Fig. 3**) shows substantial variation among brain, heart, kidney, and liver. The Venn diagram in **Figure 3a** quantifies the distributions of these adducts among the tissues, revealing that the epigenetic marks (5-MdC, 5-HMdC, 5-FdC) were detected in every tissue, as expected. While there were a few putative adducts uniquely detected in each tissue, 60-70% of the signals were detected in all tissues (52 of 92 in group #1 and 68 of 94 in group #2) (**Supplementary Table 1**). Among the adducts shared in all tissues, a principal components analysis (PCA) revealed strong tissue-specific variations in the levels of the shared adducts (**Extended Data Fig. 4a,c**), with a Bayesian statistical approach (**Extended Data Figure 4b,e**) revealing 5 clusters common to both groups of rats (**Extended Data Figure 4c,f**). This similar clustering for the two independent adductomics experiments underscores the potential biological meaning of the datasets. A further Gaussian mixture model clustering of the 52 shared putative DNA adducts from group #1 revealed that signals from heart tissue were uniquely segregated into two clusters, with Cluster III being heavily biased to sex (5 female) and age (≥16 mo) (**Fig. 3b**). A significant number of sex-biased DNA adducts and modifications were observed in the four tissues from group #1, as shown in **Figure 3c-f** and **Supplementary Table 2** (brain, 1; heart, 30; kidney, 34; and liver, 2; all with FDR = 1%). Among the epigenetic marks, significant sex differences were observed for 5-HMdC in heart and kidney tissues, and for 5-FdC in kidney, with no difference observed for 5-MdC in any tissue (**Supplementary Table 2**). While clusters were strongly homogenous in group #1 rats, it was not possible to analyze group #2 rats for sex differences since there were no females in 2 of 4 groups (**Extended Data Figure 3f-i**).

**Figure 3.**
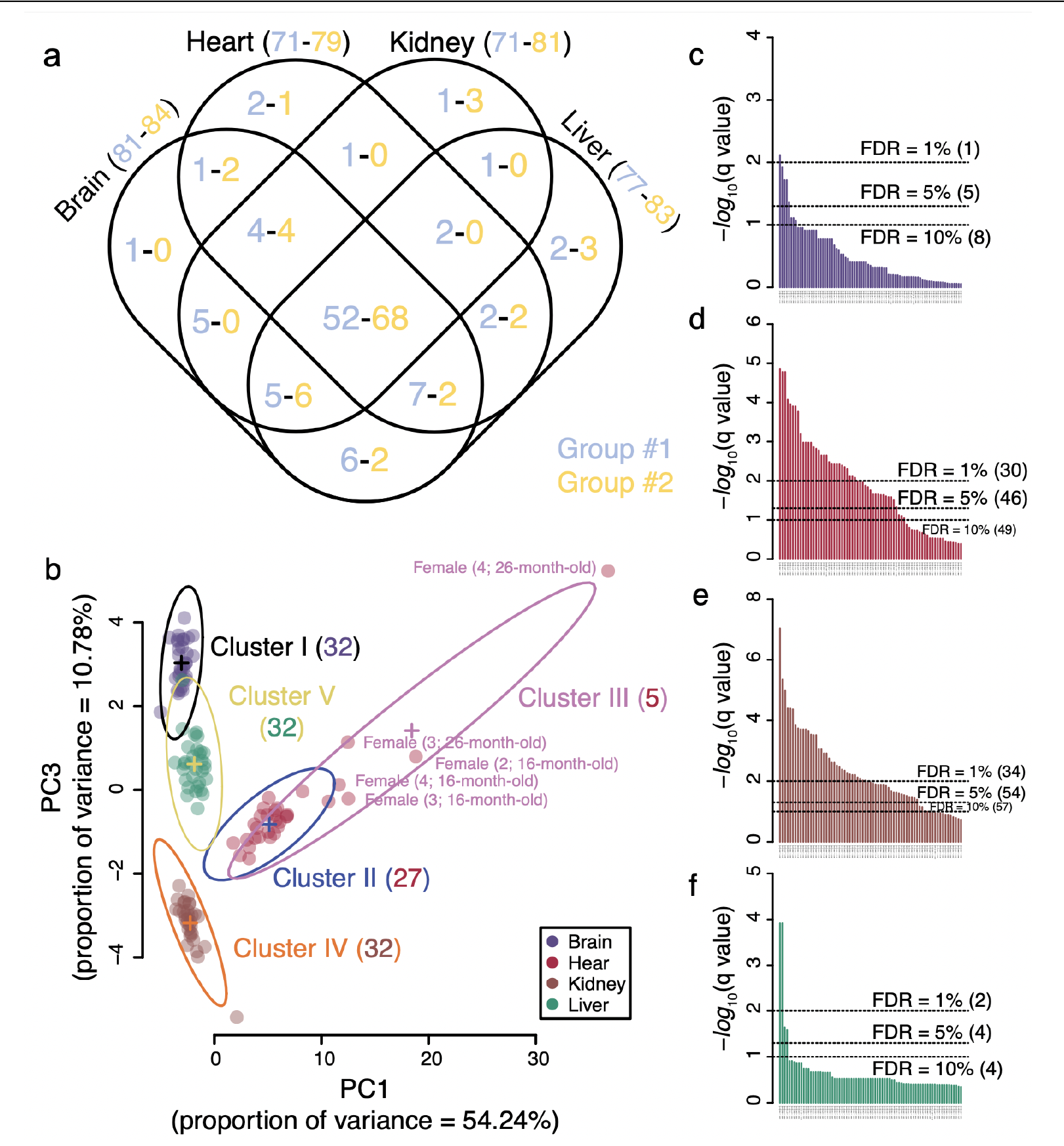
DNA damage is strongly tissue- and gender-specific in rats. (**a**) Venn diagram of the putative DNA adducts shared among the four rat tissues. Blue and yellow numbers (X-Y) correspond to the 92 putative adducts identified in group #1 rats and 94 in group #2 rats. Data are from **Supplementary Table 1**. (**b**) Based on multivariate statistical analysis of tissue, sex, and age associations of the 52 shared adducts in group #1 rats (**Extended Data Fig. 4a-c**), an EVV model (ellipsoidal distributions with equal volume and variable shape and orientation axes) revealed 5 underlying Gaussian distributions (Clusters I-V). The number of members in each cluster is shown in parentheses. Members of Cluster III comprise a striking group of 5 heart samples from mainly older female rats (labeled with sample # and age). (**c-f**) Bar plots of q-values for comparisons of putative DNA adduct levels in tissues from female versus male group #1 rats: (**c**) brain (81 adducts), (**d**) heart (71 adducts), (**e**) kidney (71), and (**f**) liver (77). q-Values were calculated from p-values obtained with the Wilcoxon rank sum test pn data from **Supplementary Tables 1** and **2**. Dotted lines represent the false discovery rates (FDR) at 10, 5, and 1%, with values in parentheses representing the number of significant adducts at each threshold.

**Figure 4.**
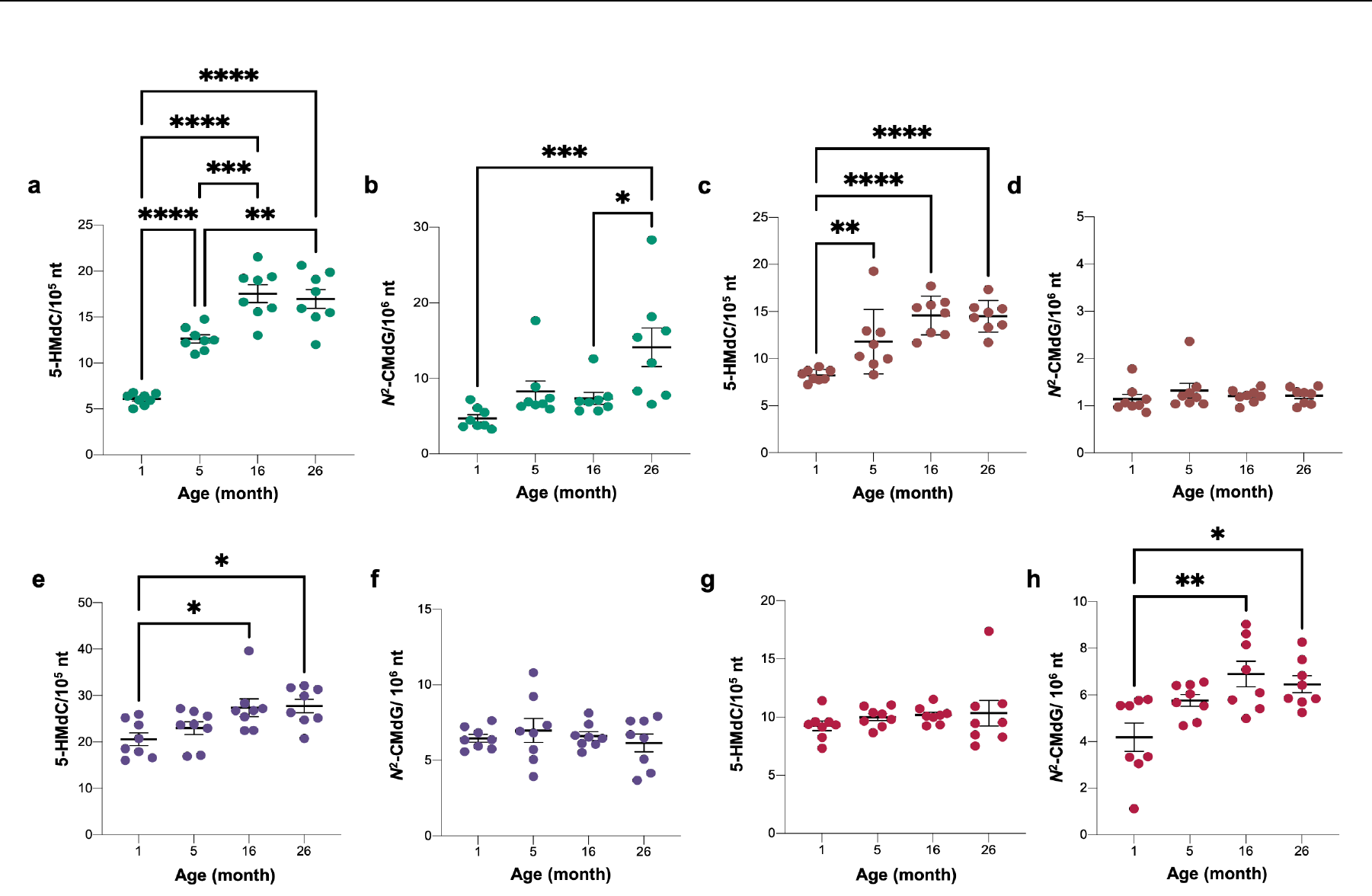
Quantitative analysis of age-dependent DNA damage products in rat tissues. Isotope dilution chromatography-coupled triple quadrupole mass spectrometric analysis was used to quantify 5-HMdC. (**a, c, e, g**) and *N*^2^-CMdG (**b, d, f, h**) levels in liver (**a, b**), kidney **(c, d**), brain (**e, f**), and heart (**g, h**) from rats at 1-, 5-, 16- and 26-months-old. Box and whisker plots show mean ± SEM for N=8. One-way ANOVA with Bonferroni’s multiple comparisons test was used to evaluate differences among the ages. \**p*<0.05, \*\**p*<0.01, \*\*\**p*<0.001 and **** *p*<0.0001 indicate statistically significant differences.

### Adductome analysis in rat tissues reveals age-dependent DNA damage and modification

While most DNA adducts in rat tissues were unchanged with age, several adducts showed strong age dependence, including 5-HMdC, *N*^2^-CMdG, *N*^2^-methyl-dG, 8-Oxo-dG, and 1,*N*^6^- εdA (**Extended Data Fig. 5, Supplementary Table 3**). To validate the adductomic results, we performed sensitive and specific absolute quantification of these adducts, using isotope dilution triple-quadrupole mass spectrometry, as shown in **Figure 4**, **Extended Data Figure 6**, and **Supplementary Table 3**. The epigenetic mark 5-HMdC was previously observed to increase with age in mouse liver, kidney and brain^16, 17^ and in human tissues^19^. Our study confirmed this increase in rats aged 1 to 26 months in liver (2.8-fold, **Fig. 4a**), kidney (1.7- fold, **Fig. 4c**), and brain (1.4-fold, **Fig. 4e**), while it was unchanged in heart (*p*= 0.62) (**Fig. 4g**). Similarly, we confirmed a previous observation of age-dependent increases in 8-Oxo-dG in rat liver^20^, with a 1.8-fold increase over 26 months of age (*p*=0.0001; **Extended Data Fig. 6e**). These results validate the quantitative robustness of the adductomic analyses. Similar analyses were not performed on the group #2.

**Figure 5.**
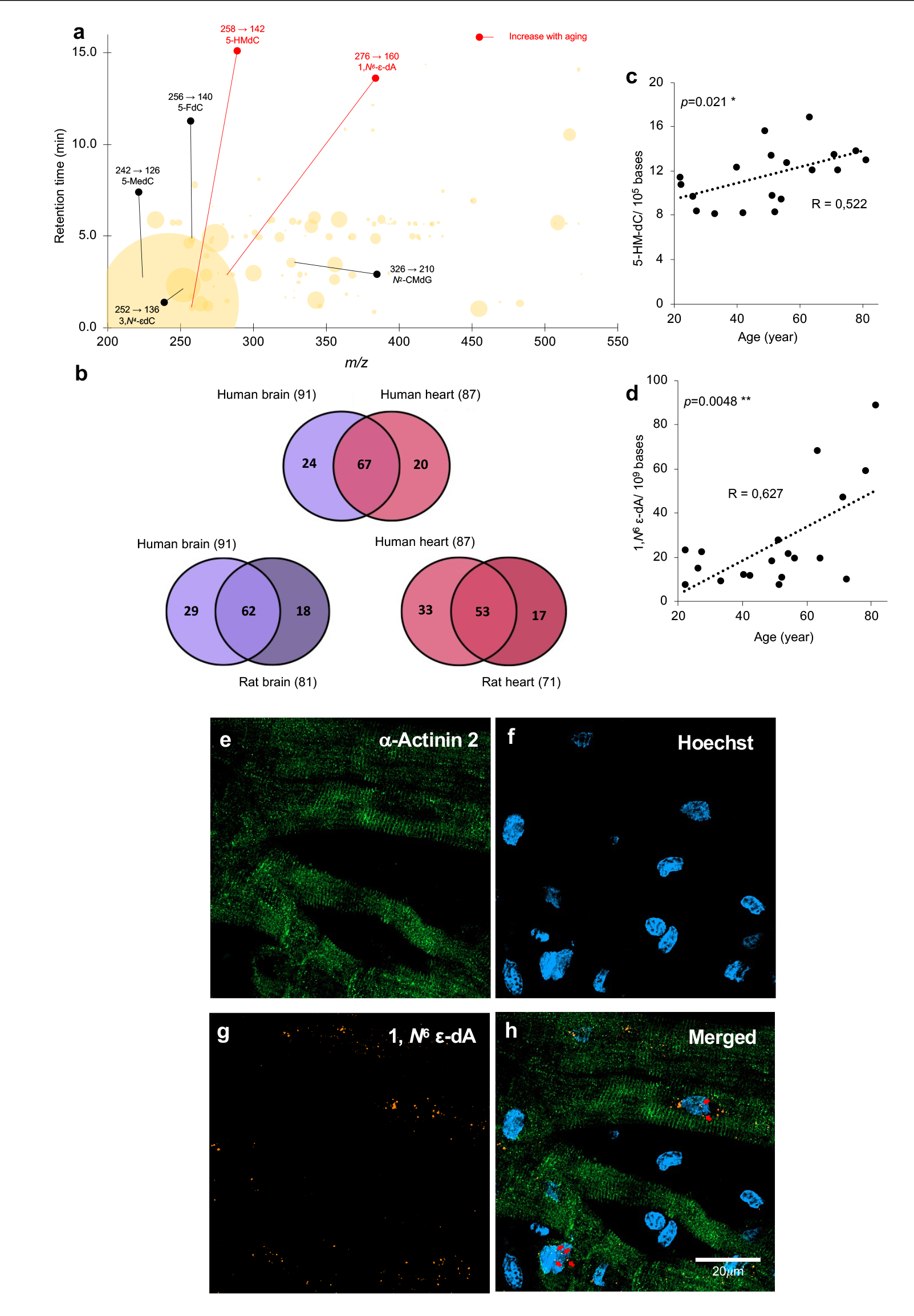
Discovery of age-tracking DNA adducts in human heart. (**a**) The bubble plot shows the DNA adductome map in non-failing human heart. Mass spectrometry transitions are noted above the names of adducts verified by chemical standards. Adducts that increase with age are highlighted in red. (**b**) Venn diagrams showing overlap among the adductomic signals detected in human and rat heart and brain. (**c,d**) Levels of 5-HMdC and 1, *N*^6^ ε-dA, 5- HMdC were quantified in 19 human non-failing heart samples from donors ranging in age from 22- to 81-years-old by isotope dilution chromatography-coupled triple quadrupole mass spectrometric analysis. Pearson’s correlation test was used to establish correlation between 5- HMdC and 1,*N*^6^-εdA levels and age; \**p*<0.05 and \*\**p*<0.01. (**e-h**) Confocal imaging of sectioned and immune-stained human non-failing heart samples reveals the presence of scattered and sparse 1,*N*^6^-εdA throughout cardiac tissue. 1,*N*^6^-εdA is present within cardiac cells, as seen by orange spots found within green (*α*-Actinin 2 labeled) areas. Red arrows denote 1,*N*^6^-εdA spots within nuclei (labeled with Hoechst), as shown in the merged figure panel (**h**). Scale bar represents a length of 20 *μ*m.

We also found age-dependent increases in *N*^2^-CMdG, an advanced glycation end-product (AGE) that results from a reaction between glyoxal and 2’-deoxyguanosine (dG)^21^ and that has been in mouse hepatoma cells, 293T human kidney cells, and calf thymus DNA^12, 22^. In rat, *N*^2^-CMdG increased in 3-fold in liver (**Fig. 4b**) and 1.5-fold in heart (**Fig. 4h**) over 1-26 months but remained unchanged in kidney (**Fig. 4d**) and brain (**Fig. 4f**). While mutations caused by glyoxal-induced DNA adducts can be prevented by nucleotide excision DNA repair (NER), mismatch DNA repair (MMR), and the protein DJ-1 *in vitro*^23, 24^, metabolically- derived glyoxal, methylglyoxal, and other reactive aldehydes are removed by the glyoxalase system of GLO1 and GLO2 in humans to prevent DNA adduct formation. In this system, GLO1 converts glyoxal, methylglyoxal, and glutathione (GSH) to *S*-glycolylglutathione that is cleaved by GLO2 to glycolate, D-lactate, and GSH^25^. Here, we quantified the effect of GLO1 expression on *N*^2^-CMdG levels in brain, heart, and liver tissues from mice with *Glo1* knock-down^26^ or over-expression^27^ and wild-type littermates. Surprisingly, the only change observed in *N*^2^-CMdG levels involved a significant increase in liver in GLO1 over-expressing mice compared to wild-type (**Extended Data Fig. 6f-j**). These data indicate either that *N*^2^- CMdG is formed from sources other than glyoxal or that GLO1 does not significantly affect glyoxal levels in mice, either of which has implications for the observed age-dependent accumulation of *N*^2^-CMdG.

### HMdC and 1,N^6^-εdA are age-dependent DNA adducts in human heart

The observed results in rats raised the question of age-dependent DNA damage and modification in humans. To this end, we performed adductomic analyses with DNA from non-failing human myocardial tissues comprised of left ventricular myocardial samples from 17 people ranging in age from 22- to 81-years-old^27, 28^ and from 10 human brain tissue samples from persons without cognitive impairment ranging in age from 71- to 89-years-old (**Supplemental Table 5**). The analysis revealed 111 total putative DNA adducts and modifications, with 91 detected in brain and 87 in heart (**Fig. 5**, **Supplementary Table 3**). The Venn diagrams in **Figure 5b** reveal that rats and humans share signals for 62 putative adducts in brain and 53 in heart.

Among these, adductomics revealed that 5-HMdC and 1,*N*^6^-εdA all increased with age in human hearts (**Fig. 5**, **Supplementary Table 3**), with this age dependence corroborated by isotope dilution LC-MS/MS that shows a significant and positive correlation for 5-HMdC (*p*= 0.021) and 1,*N*^6^-εdA (*p*= 0.0048) in non-failing human hearts (**Fig. 5c,d**; **Supplementary Table 3**). As a product of reaction of dA with electrophilic products of lipid peroxidation^29^, 1,*N*^6^-εdA levels are elevated in human diseases such as Wilson’s disease, hemochromatosis, and cirrhosis^30^, and show age-dependent increases in liver and brain in the OXYS rat model of aging due to oxidative stress^31^. **Figure 5e-h** shows 1, *N*^6^ ε-dA concentrated mainly in select cardiomyocyte nuclei, raising the question of the age of these cells in the local tissue environment.

## Discussion

In spite of the overwhelming evidence that imbalances in hundreds of types of DNA damage and dozens of DNA repair enzymes cause and drive many human diseases, such as in cancer^32^, the underlying mechanisms that lead to age-, tissue-, and sex-specificity of genetic toxicity are poorly understood. Here, we took a first step in unraveling this problem with a systems-level estimate of the spectrum of DNA damage products and modification in rat and human tissues. Such an analysis is possible only with the emergence of convergent adductomic technology consisting of untargeted mass spectrometric detection and quantification of virtually any type of nucleobase damage in DNA^6^ coupled with statistical modeling to visualize patterns and covariations among the putative DNA damage products. This systems-level untargeted DNA adductomic approach serves as a discovery tool by revealing all detectable forms of DNA damage, including new chemical species, followed by providing relative quantification of adduct levels across different conditions.

### Validation of the adductomic method

As an untargeted method, there is always concern about detecting artifacts caused by non-DNA contaminants that produce a low-resolution CID fragmentation mimicking loss of 2’-deoxyribose. Such artifacts are mitigated by starting with highly purified DNA and accounting for contaminants in non-DNA components of the analyzed samples. Each analytical run must include controls lacking buffer, enzymes, and DNA to identify contaminants. Our adductomic method helped us to achieves these requirements. Selected signals were reproducibly present at 3-times the background level in all 32 samples of each analyzed rat tissue (*i.e.,* 8 samples of each tissue at 4 different ages; 22% average error), with more than half of the detected adducts being shared between rats and humans in brain and heart tissues (**Fig. 5b**) in two different sets of rat. The adductomic method was validated (1) by *post hoc* identification of 10 detected adducts using chemical standards and (2) by high-resolution structural analysis of products, such as the tentatively identified cyclobutane dimer of dC. Of the remaining 114 unidentified species, about half had low-resolution *m/z* values consistent with known DNA damage structures (**Supplementary Table 1**). More interesting after filtering our data, we were able to find 63% of shared adducts between the two studies showing the robustness of our approach. Thus, there is a high probability that most signals detected in our adductomic analyses are indeed DNA adducts that merit identification and definition of formation and repair mechanisms to understand their roles in pathobiology and the basis for our observed age, tissue, and sex biases. Of course, the adductomic method is not as sensitive as isotope-dilution LC-MS/MS and will not detect labile nucleobase species, such as rapid depurination of 8-nitro-2’-deoxyguanosine arising during inflammatory stresses^2^, or species arising from oxidation of the 2’-deoxyribose moiety^33^. There are also complexities in the adductomic method, such as MS ionization of adducts as salts (*e.g.,* possible ammonium salt of 5-Cl-dC for *m/z* 279-163; **Supplementary Table 1**) and rearrangements of free 2’-deoxyribonuclesides (*e.g.,* sugar tautomers unique to FAPy-dG; *vide infra*). Despite these constraints, the strength of the adductomic approach lies in the insights gained from mining the systems-level datasets for correlative patterns that reveal behaviors of classes and groups of DNA adducts across different conditions of health and disease. By first identifying adductomic patterns, then identifying the chemical species driving the patterns, and finally doing the biochemical detective work, we can define the DNA damage products that truly drive human disease.

### Mining adductomic data for chemical classes of DNA adducts

There are many ways to mine adductomic datasets. One initial approach is to identify a putative DNA damage species that correlates with a condition of interest, such as the age-dependent increases in 8-Oxo-dG (**Extended Data Fig. 6e**), and then explore other mechanistically related species, such as the large and well-studied class of purine damage products arising during inflammation and other oxidative and nitrosative stresses (**Fig. 2**)^2^. Taking this path, we identified 8-Oxo-dG among the 114 adducts using a standard but failed to detect signals accordant with 8-Oxo-dA (*m/z* 268→152) or FAPy-dA (*m/z* 270→154). This is consistent with the observation of 10-fold lower levels of 8-Oxo-dA than 8-Oxo-dG and the relative amounts of 8-Oxo-dA, 8-Oxo-dG, FAPy-dA, and FAPy-dG in cells and tissues subjected to irradiation and other oxidative stresses (**Fig. 2**)^34–36^. While we did detect two signals suggestive of the known anomeric or sugar tautomer forms of FAPy-dG (*m/z* 286-170)^37–39^, the observed HPLC retention times of 2.9 and 4.9 min for *m/z* 286-170 (**Supplementary Table 1**) are inconsistent with expectations-based retention times of ∼0.7 and ∼0.9 min for 5-methyl-FAPy-dG (**Fig. 2**). We also observed signals suggestive of the other oxidized purine nucleobase products noted in **Figure 2**: oxazolone (Oz; *m/z* 247→131), nitroimidazole (NitroIm; *m/z* 287→171), spiroiminodihydantoin (Sp; *m/z* 300→184; same as FAPy-dG), and guanidinohydantoin (Gh; *m/z* 274→158). While 8-Oxo-dG and putative Oz were variably present in rat and human tissues, the putative NitroIm signal was only present in rat liver tissue and a signal consistent with Gh was present in all rat and human tissues at high levels. The observation with Gh runs contrary to published *in vitro* studies of DNA damage caused by a variety of oxidants in which Gh rose at levels nearly two orders-of-magnitude lower than 8-Oxo-dG in all cases^40^. However, the biochemical environment in cells and tissues has the potential to strongly influence the chemical reactions involved in DNA damage formation and partitioning, which leaves the door open to the identity of *m/z* 274→158 as Gh. Based on the HPLC retention times of chemical standards, we ruled out Sp and 5-methyl-FAPy-dG as the adduct present at *m/z* 300→184: ∼0.95 min for Sp (consistent with its hydrophilicity) and ∼0.7 and ∼0.9 min for the anomeric or tautomeric pair of FAPy-dG species (**Fig. 2**). The adductomic data thus provides the foundation for a host of testable hypotheses about the *in vivo* behavior of entire classes of adducts arising by similar chemical mechanisms: Are *in vitro* observations recapitulated *in vivo*? How does the cellular environment alter chemical partitioning? What are the true oxidants causing damage to dG and dA *in vivo*? Biochemical detective work, structural validation, and subsequent targeted analyses can then be used to test these models. Structural validation using high-resolution mass spectrometry and chemical synthesis is well-practiced and entirely feasible, as we demonstrated with the cyclobutane dimer of dC and *N^2^*- CMdG. One can also define the role of specific DNA repair mechanisms by comparing adductomic profiles between wild-type and DNA repair mutant animal models or across human populations with defined inter-individual repair capacities.

### DNA adduct biases among tissues

The power of comparing adductomic profiles across different conditions is illustrated by our observations of tissue-, age- and sex-specific biases in adduct spectra. Strong tissue-specific biases were observed in rat for 114 putative and validated DNA adducts (**Fig. 3; Extended Data Figures 3, 4d-f**), with a variety of tissue-specific or at least tissue-enriched DNA damage products (**Supplementary Table 1**). The normalized signal intensities for adducts noted in **Supplementary Table 1** represent quantitative metrics that allow relative comparisons of adduct levels across tissues given the precision of the LC-MS signals, with appropriate caution that the signal intensities do not reflect absolute amounts of the putative adducts due to differences in ionization and detection efficiencies. After removing the large signal for epigenetic mark 5-MdC, summing the signal values in a tissue represents a crude measure of the DNA adduct load in the tissue (number of adduct types and percentage of total adduct load in parentheses): rat brain, 3517 (81; 28%); rat heart, 1458 (71; 12%); rat kidney, 3574 (71; 29%); rat liver, 3790 (77; 31%); human brain, 6142 (91; 63%); human heart, 3584 (87; 37%). The DNA adduct loads were higher in the tissues analyzed in group #2: rat brain, 9483 (85; 24%); rat heart, 7085 (84; 18%); rat kidney, 10552 (90; 26%); rat liver, 13080 (90; 32%). The higher normalized signal intensities observed for the group #2 rat tissues compared to group #1 reflect higher sensitivity of the Agilent 6495C MS used analyze group #2 compared to the Agilent 6490 MS used for group #1. While there is little meaningful information that can be derived from the adduct load data, it is notable that both rat and human heart have the lowest level of adducts relative to other tissues and that the two rat groups shared a similar order of adduct loads (heart<brain<kidney<liver). The complexity of the adductomic datasets, as mixtures of potentially all classes of lesions type and repair, precludes comparisons to published studies showing tissue-specific variation in adduct repair for one or a few representative adducts or a single repair class^41–43^. For example, Gosponodov *et al.* estimated NER rates among rat kidney (0.1 lesions/kb/hr), liver (0.1 lesions/kb/hr), brain (0.08 lesions/kb/hr), and heart (0.04 lesions/kb/hr)^44^, while Langie *et al.* found lower BER activity in mouse brain compared to liver^45^. Cell replication rate could also account for tissue differences. For example, repair of 8-Oxo-dG and FAPy-dG occurs by OGG1 base-excision repair throughout the cell cycle and by MUTYH removal of A at 8-Oxo-dG:A mispairs during DNA replication^46^. MUTYH activity is highest in rapidly dividing cells (*e.g*., gastrointestinal tract) due to replication-coupled repair of 8-Oxo-dG:A mismatches. However, in addition to other repair metrics, there is no correlation between tissue-specific levels or loads of adducts and cell turnover rates in humans: hepatocyte cells (200-400 d)^47^, endothelial kidney cells (>1 y)^48^, cardiomyocytes (∼50 y)^49^ and neurons (lifespan)^50^. Even less is known about tissue-specific metabolism that causes DNA adducts. For example, mitochondrial density and energy activity correlate with production of reactive oxygen species and associated oxidative damage to lipids and carbohydrates, to form DNA-reactive alkylating agents, as well as to DNA and RNA directly^41, 42^. Brain and liver are lower in both metrics compared to heart and kidney^41, 42^, but, again, differences among the tissues in adduct load and the number of types of adducts in these tissues do not correlate with these metrics. These attempts to explain the observed biased tissue distribution of known and putative DNA adducts point to potential value in performing adductomic analyses in genetically engineered mouse models for the various DNA repair pathways, by identifying true substrates of DNA repair enzymes, quantifying the contribution of repair to adduct levels in each tissue, and to the age- and sex-dependence of adduct levels.

### DNA adduct biases among different cell types in tissues

One of the limitations of performing adductomic analyses on whole tissues is the potential for biases among specific cell types comprising the tissue. Enzymatic disaggregation of tissues may be the optimal approach for isolating cell populations, with the potential for adequate DNA for adductomic analyses, though DNA damage artifacts during processing are likely. Heat-induced artifacts are highly likely to occur with laser-capture microdissection, with DNA yield (30 ng/mm^2^) and low resolution also limiting its utility^51^. While the 5-10 μm resolution of imaging mass spectrometry may provide adequate resolution in tissues, the method is likely to be limiting in terms of sensitivity (1 fmol)^52^. Immunohistochemistry with highly specific antibodies ultimately optimizes chemical specificity with resolution. Here we used immunostaining to detect 1,*N*^6^-εdA in the nuclei of cardiomyocytes in human ventricular myocardial samples, which indicated that accumulation may be limited to a small number of cells. While this may result from a sensitivity threshold and all cells show some level of the adduct, substantial accumulation of DNA adducts in a small number of cells might indicate damage specific to cell subtypes or to the oldest cells in the tissue, with the latter indicating compromised genomic integrity during senescence.

### Age dependence of DNA adducts

Comparative analysis of DNA adducts in rats across the age span of 1 to 26 months revealed that a subset of DNA adducts did not change, several adducts showed significant age-dependent accumulation and decreases with age in a tissue- and sex-specific manner (**Fig. 4**, **Extended Data Fig. 6**). This suggests shifts in the balance between adduct formation and repair mechanisms over the life of the animal. We observed significant age-dependent increases of *N*^2^-CMdG in rat liver and heart (**Fig. 4**). *N*^2^-CMdG and the related *N*^2^-(1-carboxyethyl)-2’-deoxyguanosine (*N*^2^-CEdG) are advanced glycation end-products resulting from the reaction of glyoxal and methylglyoxal, respectively, with 2’-deoxyguanosine in DNA. This is not surprising since the methylglyoxal and glyoxal are reactive α-oxoaldehydes formed in abundance during the glucose metabolism^53^ in liver^54^ and there is evidence for age-dependent decreases in base excision repair (BER) mechanism in mouse liver^55^, with *N^2^*-CMdG likely repaired by the BER enzyme alkyladenine glycosylase (Aag)^56^. In studies examining age-dependent mechanisms of formation, we observed no effect of glyoxalase I activity on *N*^2^-CMdG levels. It is known that glyoxalase I prefers methylglyoxal as a substrate over glyoxal and protects against *N*^2^-CEdG^57, 58^. Interestingly, we detected a signal consistent with *N*^2^-CEdG at *m/z* 340→224 and found strong covariance of this signal with that for *N*^2^-CMdG (**Extended Data Fig. 6k, Supplementary Table 1**).

Adductomic analysis of human heart tissue also revealed age-dependent DNA damage and epigenetic marks. We observed a significant age-dependent accumulation of 5-HMdC, which results from oxidation of 5-MdC^19^, in non-failing heart tissue, which parallels a published observation of increases in 5-HMdC with age in grey and white matter of the human cerebrum^17^. We also found age-dependent increase in the 1,*N*^6^-εdA damage product. While this contrasts the inconsistent changes in 1,*N*^6^-εdA in rat heart (**Extended Data Fig. 6d**), it is consistent with the idea of cardiomyocytes becoming more susceptible to oxidative stress with age^59^, with cardiac DNA repair mechanisms tending to be less efficient with age^60^, and with the observation of age-dependent increases in the 4-hydroxy-2-nonenal (HNE) lipid peroxidation product that reacts with DNA to form 1,*N*^6^-εdA^61, 62^. The immunohistochemical evidence for concentration of 1,*N*^6^-εdA in subpopulations of cardiomyocyte nuclei suggests a possible correlation between adduct formation and cell age or senescence.

### Sex dependence of DNA adducts

The complexity of the network of determinants of DNA adduct spectra is increased by our observation of sex-specific differences in adduct levels in all four of the tissues studied (**Fig. 3c-f, Supplementary Table 2**). Interestingly, while the epigenetic mark 5-MdC did not show a sex bias in any of the rat tissues, its oxidation products 5-HMdC and 5-FdC were significantly higher or trending higher in female kidney and heart (**Supplementary Table 2**). While sex-based biases in metabolism may account for the observation that females show higher levels of smoking-related DNA adducts in human lung tissue^63^ and in mice treated with 2-amino-3-methylimidazo [4,5-f]-quinoline, a potent food mutagen^64^, our observations with 5-HMdC and 5-FdC suggest sex differences in epigenetic regulation of gene expression. The variety of unknown putative DNA adducts showing sex biases merits further investigation of their structures and definition of their potential roles in cell biology, aging, and pathology.

## Conclusions

Adductomics proved to be an effective approach for discovering individual DNA damage products as well as for systems-level analysis of the behaviors of dozens of putative DNA adducts as a function of tissue type, age, and sex. Such covariate behavior can be highly informative about local environment of DNA repair and metabolism, especially when combined with large metabolomics^65^, proteomics^66, 67^, and other datasets. While age- sex-, and tissue-specific variations in the levels of 114 putative and method-validated DNA adducts provide important new biological insights into genetic toxicology and epigenetics, the adductomic approach illustrated here is a powerful approach to (1) assigning substrates to DNA repair pathways using genetically engineered animal models, (2) understanding mechanisms of DNA damage formation by monitoring classes of DNA lesions and manipulating metabolic pathways, and (3) quantifying inter-individual variation in DNA damage and repair across populations and pathologies. These efforts will be enhanced by advancing the mass spectrometry and data analysis tools for adductomic profiling^67^.

## Supporting information

endnot library

## Methods

### Materials

Tris-hydrochloride (Tris HCl), magnesium chloride (MgCl2), acetic acid, sodium acetate, formic acid, butylated hydroxytoluene (BHT), tetrahydrouridine (THU), coformycin, DNAse I, benzonase, phosphatase alkaline, desferrioxamine mesylate salt, glyoxal, 4-Thio-2’- deoxyuridine, 6-thio-2’-deoxyguanosine, 5-Methyl-2’-deoxycytidine, 2-Chloro-2’- deoxyadenosine, 8-Hydroxy-2’-deoxyguanosine, 5-hydroxymethyl-2’-deoxyuridine, 5-Fluoro- 2’-deoxycytidine and 6-thio-2’-deoxyguanosine were obtained from Sigma-Aldrich (Saint- Louis, MO, USA). Phosphodiesterase I was purchased from Affymetrix (Cleveland, Ohio, USA). [^15^N(U)]-labeled 2’-deoxyguanosine was purchased from Cambridge Isotopes (Andover, MA, USA). Isopropanol and acetonitrile were purchased from VWR Scientific (Franklin, MA, USA). Deionized water was further filtered through MilliQ systems (Millipore corporation, Bedford, MA, USA) and used in the whole experiment. Anti-sarcomeric protein α-Actinin 2 and ProLong Gold Antifade were purchased from Thermofisher Scientific (Waltham, MA, USA) while 1, *N*^6^ ε-dA Antibody and TrueBlack Plus Lipofuscin Autofluorescence Quencher came from Novus Biological (Centennial, CO, USA) and Biotium (Fremont, CA, USA) respectively. 5-Hydroxymethyl-2’-deoxycytosine, 1, *N*^6^ ε-2’- deoxyadenosine, 5-formyl-2’-deoxycytidine, 7-deaza-2’-deoxyxanthosine, 2’-deoxycytidine-5- carboxylic acid sodium salt, *O*^6^-nethyl-2’-deoxyguanosine, 8-oxo-2’-deoxyadenosine, *N*^2^- methyl-2’-deoxyguanosine and *N*^6^-methyl-2’-deoxyadenosine were purchased from Berry & Associates. 2’-Deoxyxanthosine, 8-chloro-2’-deoxyadenosine and 8-chloro-2’- deoxyguanosine were purchased from Biosynth Carbosynth (San Diego, CA, USA). [^3^D]-5- HMdC, [^15^N5]-1, *N*^6^ ε-dA, and [^15^N5]-8-oxo-2’-deoxyguanosine were purchased from Toronto Research Chemical (Toronto, Canada). 5’-Ethynyl-2’-deoxycytidine and 5’-chloro-2’- deoxycytidine were obtained from Cayman Chemical Company (Ann Arbor, MI, USA) and Biolog (Hayward, CA, USA), respectively.

### Synthesis of N^2^-CMdG and [^15^N(U)]-N^2^-CMdG

For unlabeled *N*^2^-CMdG, 1 mmol (0.267 g) of 2’-deoxyguanosine or [^15^N(U)]-labeled 2’-deoxyguanosine was reacted with 5 mmol of glyoxal (0.290 g) in 1 M NaOAc/HOAc buffer pH 5.5 and the mixture was heated at 37 °C overnight. The final *N*^2^-CMdG product was purified from this mixture by HPLC with a Hypersil GOLD C18 column (3.0 *×* 250 mm, 5 μm, Thermo Scientific) on an Agilent 1200 HPLC system. Binary mobile phase flow rate was 1 mL/min (A – 5mM ammonium acetate in water, B – acetonitrile; increased to 40% of B for 20 min; increased to 90% of B in 0.1 min and hold at 90% of B for 10 min; decreased to 0% of B in 1 min and re-equilibrate for 10 min). UV (*λ*max, pH 7) = 260 nm. [M+H] ^+^obs = 326.1103, [M+H] ^+^calc = 326.1111, error = 2.39 ppm. MS/MS (ESI+): 210 (nucleobase + H), 164 (nucleobase – CO2 + H), 117 (oxonium ion of 2’-dR).

### Animal procedures

All animal experiments were carried out by the local animal care and use committee and respected the principals of animal experimentation (protocol number: 16-05- 271, National Institute on Aging). For the aging study, Brown Norway wild-type rats at 1, 5, 16 and 26 months old (4 females and 4 males per group of age) or 4, 9, 17 and 26 months old (4 females and 4 males for groups at 9 and 17 months old; 8 males for groups at 4 and 26 months old) were purchased from the National Institute of Aging colony housed at Charles River. All rats had *ad libidum* access to a standard chow diet (irradiated LabDiet Prolab Isopro RMH 3000 5P75; LabDiet, St. Louis, MO) and water. All animals were maintained under a 12-hour light/dark cycle at 22 °C. After two days acclimatization, all rats were euthanized using isoflurane and cervical spine dislocation. Liver, kidneys, brain, and heart were removed, washed with ice-cold PBS, and immediately plunged into liquid nitrogen and stored at -80 °C.

For the transgenic mice, two models were used.: Reduced GLO1 activity in liver (n=11 and 18), brain (n=10 and 15) and heart (n=8 and 12) was modeled using Glyoxalase I Knock Down (Glo1-KD) mice on a C57/BL6 background and their wild-type littermates^26^. Increased GLO1 in brain (n=5 and 5) and heart (n=7 and 7) was modeled using Glyoxalase I over- expression mice on a C57/BL6 background and their wild-type littermates^27^. Frozen tissues were directly obtained from University of California San Diego. The ages for the knockdown and the overexpressing mice were 217-273 and 180-292 days, respectively.

### Human samples

Procurement of 19 frozen samples of non-failing left ventricular myocardial tissue from human donors, deemed unsuitable for transplant, were obtained from University of Pennsylvania (Philadelphia, PA USA)^68^. Samples were immediately stored at -80 °C until DNA extraction. The ages of the 9 female and 10 male subjects ranged from 22 to 81 years.

Donor demographics are detailed in **Extended Data Table 1**. Samples of human brain tissue from participants in the Religious Orders Study or Rush Memory and Aging Project (ROSMAP) without cognitive impairment (control subjects, CT) ranging in age from 71 to 89 years were obtained from the Rush Alzheimer’s Disease Center (Chicago, IL, USA)^69^ and were stored at -80 °C. Both studies were approved by an Institutional Review Board of Rush University Medical Center. Each participant signed an informed consent. Anatomic Gift Act, and Repository Consent to allow their data to be repurposed. NCI was defined as previously reported^70^. Donor demographics and definitions of clinical and neuropathological criteria are detailed in **Extended Data Table 1.**

### DNA extraction, purification, and hydrolysis

DNA extraction from rat organs (livers, kidneys, brains, and hearts), mouse organs (livers, brains, and hearts) and human hearts and brains was performed with a commercial kit (Sigma-Aldrich, 11814770001) according to the manufacturer’s protocol, with the addition to all solutions of appropriate inhibitors and antioxidants to minimize adventitious damage: 5 μg/ml coformycin, 50 μg/ml tetrahydrouridine (THU), 100 μM desferrioxamine, 100 μM butylated hydroxytoluene (BHT)^16^. DNA was then quantified by spectrophotometry at 260 nm (NanoDrop Lite Spectrophotometer, Thermofisher scientific). Samples were finally stored at -80 °C.

For DNA adductome analysis, a mix of genomic DNA (containing 5 μg of each sample to cover adductome in every age) was dried by using speed vacuum and then reconstituted in 90 μL of 10 mM Tris-HCl buffer pH 8, 1 mM MgCl2, 10 μg/mL coformycin, 50 μg/mL THU, 1 mM desferrioxamine, 1 mM BHT and digested using 10 U benzonase, 5 U DNAse I, 17U alkaline phosphatase, 0.1 U phosphodiesterase I. A blank sample was also prepared using the same procedure without the DNA. The digestion was allowed to occur overnight at 37 °C. Reactions were then worked up using 10 KDa exclusion filters and centrifuged at 12,800 × g for 10 min to remove proteins and then analyzed by LC-MS/MS.

For adduct quantification analysis, 20 μg of genomic DNA was dried by using speed vacuum and then reconstituted in 27 μL containing 10 mM Tris-HCl buffer pH 8, 1 mM MgCl2, 10 μg/mL coformycin, 50 μg/mL THU, 1 mM desferrioxamine, 1 mM of BHT and digested using 10 U benzonase, 5 U DNAse I, 17 U alkaline phosphatase, 0.1 U phosphodiesterase I. An additional 3 μL of stable isotope-labeled internal standards were then added as needed, including 5 nM [^15^N5]-*N*^2^-CMdG 1 μM [D^3^]-5-HMdC and 500 pM [^15^N5]- 1, *N*^6^-ε-dA. The digestion was allowed to occur overnight at 37 °C. Reactions were then processed using 10 KDa exclusion filters and centrifuged at 12,800×g for 10 min to remove proteins, followed by analysis by LC-MS/MS.

### DNA adductome analysis: Adduct discovery in pooled DNA samples

Adductome analysis was performed on an HPLC Agilent 1290 Infinity II with an added Diode Array Detector (DAD) coupled with the Agilent 6490 (Group #1 rats, mice, humans) or 6495 (Group #2 rats) triple quadrupole mass spectrometer with an electrspray ion source operated in positive ion mode with the following source parameters: drying gas temperature 200 °C with a flow of 12 L/min, nebulizer gas pressure 30 psi, sheath gas temperature 300 °C with a flow of 12 L/min, ESI capillary voltage 4000 V and nozzle voltage 0 V. A Kinetex EVO C18 column (2.1 × 150 mm, 2.6 μm, Phenomenex) was used for chromatographic separation (column temperature 30°C). Binary mobile phase flow rate was 0.400 mL/min (A – 0.1% formic acid in water, B – 0.1% formic acid in acetonitrile; delivered at 0% of B for 2 min; increased to 16% of B for 15 min; increased to 80% of B in 1 min and hold at 80% of B for 5 min; decreased to 0% of B in 1 min and re-equilibrate for 5 min). For 5-HMdC quantification, binary mobile phase was the following: A – water, B – 5 mM of Ammonium Acetate in water, pH 5.3; delivered at 0% of B for 2 min; increased to 16% of B for 15 min; increased to 80% of B in 1 min and hold at 80% of B for 5 min; decreased to 0% of B in 1 min and re-equilibrate for 5 min). The effluent from the first min from the LC system was diverted to waste to minimize the contamination of the ESI source. The strategy was designed to detect the neutral loss of 2’-deoxyribose from positively ionized 2’-deoxynucleoside adducts by monitoring the samples transmitting their [M+H]^+^→[M+H-116]^+^ transitions. For both the mix of DNA and the blank, 300 MRM transitions were monitored from transition *m/z* 225→109 to transition *m/z* 525→409 with a collision energy of 10 eV. Ten microliters of both samples were injected six times to cover the monitoring of 300 MRM transitions. Each of 8 methods covering the 300 MRM transitions has a final cycle length/ scan time of 790.4 ms (1.27 cycles/s). Transitions of canonical 2’-deoxynucleosides, including *m/z* 228→112 ([dC+H]^+^), *m/z* 243→127 ([dT+H]^+^), *m/z* 252→136 ([dA+H]^+^) and *m/z* 268→152 ([dG+H]^+^), were not monitored in this adductome analysis. Finally, the 300 MRM signals were compared between the mix of DNA and the blank. Each signal that was present in the mix of DNA sample but not in the blank was collected for further investigation. These stepped MRM signals were normalized to account for different amounts of injected DNA with each LC-MS run by dividing the mass spectrometer signal intensity by the sum of the absorbance values from an in-line diode array UV detector for the 4 canonical 2’-deoxyribonucleosides. The resulting normalized signal intensities, detailed in **Supplementary Table 1a** for both groups of rats, were curated to remove artifacts, noise, and other spurious signals using the decision tree detailed in **Figure 1a** to remove common salts, isotopomers, known artifacts, and potential noise (signal intensity <2). The resulting relative peak intensity of these signals are detailed in **Supplementary Table 1e** and plotted as a bubble chart in which the x-axis was the *m/z* and the y-axis was retention time (**Fig. 1**).

### DNA adductome analysis: Adduct quantification in individual DNA samples

The final set of 114 putative DNA adducts and modifications detailed in **Supplementary Table 1e** was used for targeted analysis of individual tissue samples from rats. Portions of isolated DNA used for the adduct discovery described earlier were analyzed again with an MRM table for the 114 putative DNA adducts and modifications and the same LC-MS/MS conditions. The resulting signals were normalized by dividing by the sum of the UV signals for the four canonical 2’-deoxyribonucleosides, as described earlier. The normalized signal intensities for both groups of rats are detailed in **Supplementary Table 4**.

### Identification of DNA damage products with standards

To tentatively identify the structures of adductomic signals, a cocktail of 2’-deoxyribonucleoside standards (10 nM each in 30 μL of water) was analyzed by LC-MS/MS using the same conditions noted earlier for adductomic analysis. The goal here was to correlate retention times and MS/MS transitions for the standards with the adductomic data. The following transitions were monitored: *m/z* 326 → 210 *N*^2^-carboxymethyl-2’-deoxyguanosine, *N*^2^-CMdG), *m/z* 258 → 142 (5-hydroxymethyl-2’- deoxycytidine, 5-HMdC), *m/z* 276 → 160 (1,*N*^6^-etheno-2’-deoxyadenosine, 1,*N*^6^ ε-dA), *m/z* 242 → 126 (5-methyl-2’-deoxycytidine, 5-MdC), *m/z* 245 → 129 (4-thio-2’-deoxyuridine, 4- thio-dU), *m/z* 252 → 136 (3,*N*^4^-etheno-dC, 3, *N*^4^-ε-dC), *m/z* 256 → 140 (5-formyl-2’- deoxycytidine, 5-formyl-dC), *m/z* 262 → 146 (5’-chloro-2’-deoxycytidine, 5-CldC), *m/z* 268 → 152 (7-deaza-2’-deoxyxanthosine, 7-deaza-dX), *m/z* 268 → 152 (8-hydroxy-2’- deoxyadenosine, 8-Oxo-dA), *m/z* 269 → 153 (2’-deoxyxanthosine, dX), *m/z* 274 → 158 (guanidinohydantoin, Gh), *m/z* 282 → 166 (*O*^6^-methyl-2’-deoxyguanosine, *O*^6^-MdG), *m/z* 282 → 166 (*N*^2^-methyl-2’-deoxyguanosine, *N*^2^-MdG), *m/z* 284 → 168 (8-hydroxy-2’- deoxyguanosine, 8-Oxo-dG), *m/z* 284 → 168 (6-thio-2’-deoxyguanosine, S6-dG), *m/z* 294 → 178 (2’-deoxycytidine-5’-carboxylic acid sodium salt, dC carboxylic acid), *m/z* 300 → 184 (spiroiminodihydantoin, Sp), *m/z* 304 → 188 (pyrimidopurinone adduct of 2’- deoxyguanosine, M1-dG), *m/z* 286 → 170 (8-chloro-2’deoxyadenosine, 8-Cl-dA), *m/z* 302 → 186 (8-chloro-2’-deoxyguanosine, 8-Cl-dG), *m/z* 286 → 170 (2-chloro-2’-deoxyadenosine, 2- Cl-dA) and *m/z* 246 → 130 (5-fluoro-2’-deoxycytidine, 5-F-dC).

### Identification of the cyclobutene dimer of dC

The identity of the putative DNA adduct with *m/z* 455 → 339 was achieved by multi-stage mass spectrometry (MS^n^) using an HPLC- coupled Thermo Orbitrap ID-X Tribrid mass spectrometer coupled with a Vanquish LC (Thermo Fisher). Five microliters of sample were injected on a Kinetex C18 column (150 × 2.1 mm, Phenomenex) eluted with mobile phase A (0.1% formic acid) and mobile phase B (0.1% formic acid in acetonitrile) using the following gradient: 2 min at 0% then to 18% B in 13 min and to 60% B in 2 min. The original conditions were re-established in 1 min and the column re-equilibrated for 5 min. MS^1^ data were acquired at 120,000 resolution in the orbitrap with a 70% RF lens. MS^2^ data were acquired on the targeted *m/z* identified previously, using assisted HCD spanning 20, 35, and 60 normalized collision energy, and assisted CID at 15,30,45 %. MS^2^ fragments were measured in the orbitrap at 60,000 resolution. MS^2^ fragments with >5% relative intensities were selected for MS^3^, with a fragmentation by CID at 30%, and analysis in the orbitrap at 30,000 resolution. All data were analyzed in xcalibur qual browser, with some of the fragmentation additionally analyzed in Massfrontier (V8.0, Thermo fisher).

### DNA adduct quantification by isotope dilution LC-MS/MS

For absolute quantification of specific DNA modifications and damage products, isotope dilution LC-MS/MS was performed on isolated DNA used for adductomic analyses. Ten microliters of each sample were injected into the LC-MS/MS instrument using the same LC-MS/MS parameters noted in “DNA adductome analysis”. Characteristic reactions and collision energies for DNA adducts of interest are as follows (*m/z* precursor ion → *m/z* product ion, collision energy (eV), retention time): *N*^2^-CMdG (*m/z* 326→*m/z* 210, 10, 3.45min), [^15^N5]-*N*^2^-CMdG (*m/z* 331→*m/z* 215, 10, 3.45min), 5-HMdC (*m/z* 258→*m/z* 242, 10, 2.72min), [D^3^]-5-HMdC (*m/z* 261→*m/z* 245, 10, 2.71min), 1, *N*^6^-ε-dA (*m/z* 276→*m/z* 160, 10, 2.93min), [^15^N5]- 1, *N*^6^ ε-dA (*m/z* 281→*m/z* 165, 10, 2.93min), 8-Oxo-dG (*m/z* 284→*m/z* 170, 10, 6.65min), and [^13^C,^15^N2]-8- Oxo-dG (*m/z* 286→*m/z* 170, 10, 6.65min). Calibration curves for the labeled and unlabeled forms were constructed by plotting the MRM signal ratios between the labeled and unlabeled forms against their corresponding concentration ratios, as illustrated in **Extended Data Figure 1**. Quantification for *N*^2^-CMdG, 5-HMdC, 1, *N*^6^-ε-dA and 8-Oxo-dG were achieved using the ratio of peak areas of the unlabeled and labeled DNA adducts. Raw mass spectrometric data were analyzed by MassHunter Qualitative Analysis software (Agilent Technologies, USA).

### Immunostaining and imaging

To prepare samples for immunostaining, samples were dissected from patients and stored at -80 °C freezer prior to transferring to Tissue-Tek optimal cutting temperature (OCT) compound for cryosection. Slides with sectioned samples were then fixed with ice-cold 100% methanol for 15 min at 4 °C, washed 3 times with PBS, and blocked for 1 h with a 5% goat serum, 0.3% Triton X-100 and PBS solution. Primary antibodies were then applied to samples in antibody buffer (1% BSA and 0.3% Triton X-100 in PBS), overnight at 4 °C. For cardiomyocyte staining, anti-sarcomeric protein α-actinin 2 was applied at 1:250 or 2 μg/mL, and 1,*N*^6^-εdA antibody was applied at 1:100, in antibody buffer. Following five washes in PBS 0.3% Triton X-100, secondary antibodies were then applied for 1hr in antibody buffer, at 1:1000. α-Actinin 2 was detected with Goat anti-Rabbit IgG conjugated to Alexa 488, and 1,*N*^6^-ε-dA was detected with Anti-Mouse IgG conjugated to Alexa 555. To detect nuclei, Hoechst 33342 was applied for 10 min following secondary antibody removal, at 1:10,000 in a 0.3% Triton-PBS. Slides were then washed 5 times in 0.3% Triton-PBS, washed two times in PBS, and TrueBlack Plus Lipofuscin Autofluorescence Quencher was applied for 10 min to reduce autofluorescence. Finally, TrueBlack was washed 3 times with PBS, and slides were mounted in ProLong Gold Antifade. Images were captured with an LSM 980 Confocal Microscope (Zeiss), using the Airyscan mode and a 63x oil objective. Z-stacks were captured across nine tiles, to sample larger areas for image analysis.

### Image analysis

To detect 1,*N*^6^-ε-dA in cardiac cell nuclei within sectioned samples, we performed image analysis on captured confocal images using arivas Vision 4D software (arivas). We used the machine learning module to detect cardiac cell networks identified by *α*-actinin 2 staining, the thresholding segmentation module to detect nuclei by Hoechst 33342 staining, and the blob finder module to detect 1,*N*^6^-εdA spots. Then we used the compartments module to detect nuclei within cardiac cells networks and 1,*N*^6^- εdA within these nuclei. Selected nuclei were encompassed by greater than 70% of the cardiac signal, and 1,*N*^6^-εdA puncta were selected when they were encompassed by greater than 50% nuclei signal in selected nuclei.

### Statistical analyses

All statistical analyses were performed using either GraphPad Prism (version 8.0) or R (http://www.r-project.org). Data are presented as means ± SEM (standard error of the mean). Each biological sample was prepared at least two times to confirm observation. Statistical significance was determined using one-way ANOVA with Bonferroni’s multiple comparisons test for rat study and a two-sided unpaired Student’s *t*-test to compare wild-type (WT) to KD/OE mice samples. Finally, we used Pearson’s correlation in human non-failing heart samples. * *p* < 0.05, ** *p* < 0.01, *** *p* < 0.001 and **** *p* < 0.0001 was considered to indicate statistically significant differences. All statistical analyses were performed using GraphPad Prism version 8.0 (GraphPad Software).

We traced differences of DNA adducts in rat sex and age using two-sided Wilcoxon rank sum test (equivalent to the Mann-Whitney U test; wilcox.test function of stats package in R). The false discovery rate (FDR), the rate of type I error associated with rejecting the null hypothesis due to multiple comparisons, was estimated using qvalue function in the qvalue package^71^ in R.

To identify the most strongly covarying DNA adducts, we performed principal component analysis (PCA) on shared adducts in rat samples (**Figure 3a, Extended Data Figure 3c**) in both group, that is the most commonly used method for reduction of dimensionality^72^, using the prcomp function (stats package in R). To find underlying patterns, we used mixture model clustering that is a probability-based approach in which we assume the data set is best described as a mixture of probability models. We employed the mclust package^73^ for Gaussian mixture modeling (GMM), the most used mixture model-based clustering method, to determine the underlying Gaussian mixture distributions using first two PC scores covering 62,71% variation of the data (**Extended Data Fig. 4c,f**).

To assess association between 162 DNA adducts, we estimated Spearman’s rank correlation coefficient (*ρ*) after zero filling the empty cells (**Supplementary Table 2**). Results are show as a heatmap reordered by hierarchical clustering analysis (HCA) obtained by the R pvclust package^74^ after 3000 iterations of multiscale bootstrap resampling.

## Acknowledgments

The authors gratefully acknowledge support from grant R01-AG063341 (PIs: Lee and Dedon) from the National Institute on Aging of the National Institutes of Health and from Center grant P30-ES002109 from the National Institute of Environmental Health Sciences of the National Institutes of Health. ROSMAP is supported by P30AG10161, P30AG72975, R01AG15819, R01AG17917, U01AG46152, U01AG61356 (to D.A.B.). ROSMPA resources can be requested at http://www.radc.rush.edu.

## Conflict of interest

The authors have no conflicts of interest to declare.

## Author contributions

P. Dedon and R. Lee conceived of the study, supervised all aspects of the study, and interpreted data. A. Guilbaud, F. Ghanegolmohammadi, Y. Wang, A. Kreymerman, J. Gamboa-Varela, J. Leng, J. Garbern, E. Ricci-Blair, H. Elwell, C. Vidoudez, F. Cao, S. Balamkundu, M. DeMott and Cui L. conducted the experiments, analyzed, and interpreted the data. K. Bedi, K. Margulies, D. Bennett, A. Palmer and A Barkley-Levenson participated in study design and provided key biological specimens. A. Guilbaud, F. Ghanegolmohammadi, Y. Wang, and A. Kreymerman drafted the manuscript and all authors contributed to reviewing and revising the manuscript.

## Data availability statement

Mass spectrometry data that support the findings of this study have been deposited in the Chorus Project (https://chorusproject.org/pages/index.html) with the accession code 1767.

## Competing Interests Statement

A. Palmer has a patent for the methods and use of Glo1 inhibitors (https://patents.google.com/patent/US11235020B2/en).

## Corresponding Author

Correspondence and requests for materials should be addressed to Peter Dedon at.

## Supplementary Information

Supplementary Information is available for this paper:

Supplementary Table 1. DNA adduct discovery: Normalized adductomic data for age-pooled samples in rat tissues from two independent experiments (rat groups #1 and #2). Separate sheets for each step of data filtering.

Supplementary Table 2. Analysis of adductomic data in individual DNA samples in rat group #1 as a function of sex and age.

Supplementary Table 3. DNA adduct levels in mouse, rat, and human tissues measured by isotope dilution LC-MS/MS.

Supplementary Table 4. Normalized mass spectrometric signals for individual samples of rat tissues from rat groups #1 and #2.

Supplementary Table 5. DNA adduct discovery and normalized signal data for human brain and heart samples.

## Extended Data Figure Legends

**Extended Data Fig. 1.**
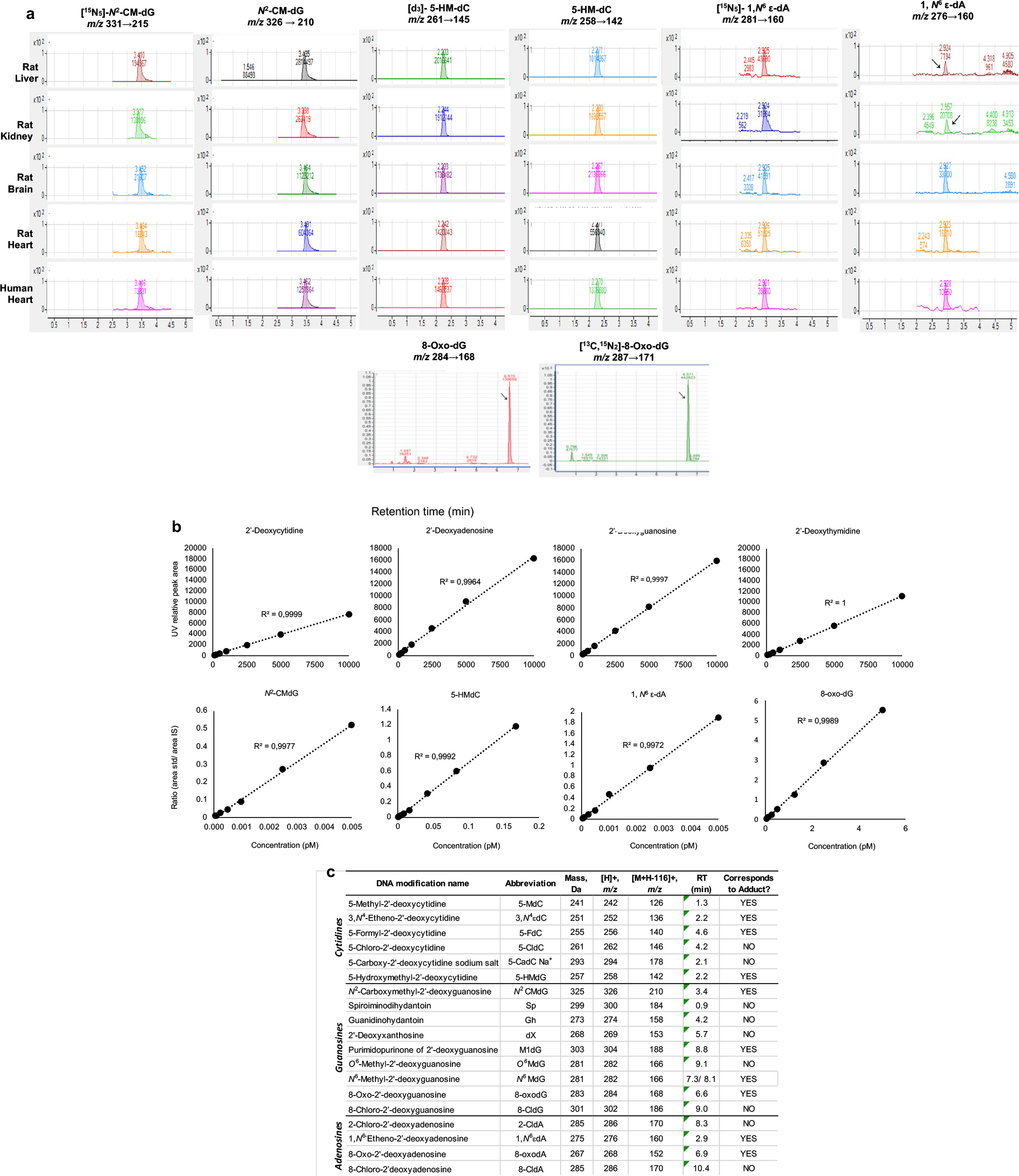
Parameters and data for isotope-dilution chromatography-coupled triple quadrupole mass spectrometric quantification of DNA adducts in rat and human tissues. (**a**) Representative extracted ion chromatograms for analysis of *N*^2^-CMdG, 5-HMdC,8-oxo-dG, and 1,*N*^6^-εdA and their isotope-labeled counterparts in rat and human tissues. (**b**) Illustrative calibration curves of the canonical 2’-deoxyribonucleosides and several DNA adducts. (**c**) LC-MS/MS parameters for known DNA adducts and modifications analyzed here. These data pertain to Figures 4 and 5.

**Extended Data Figure 2.**
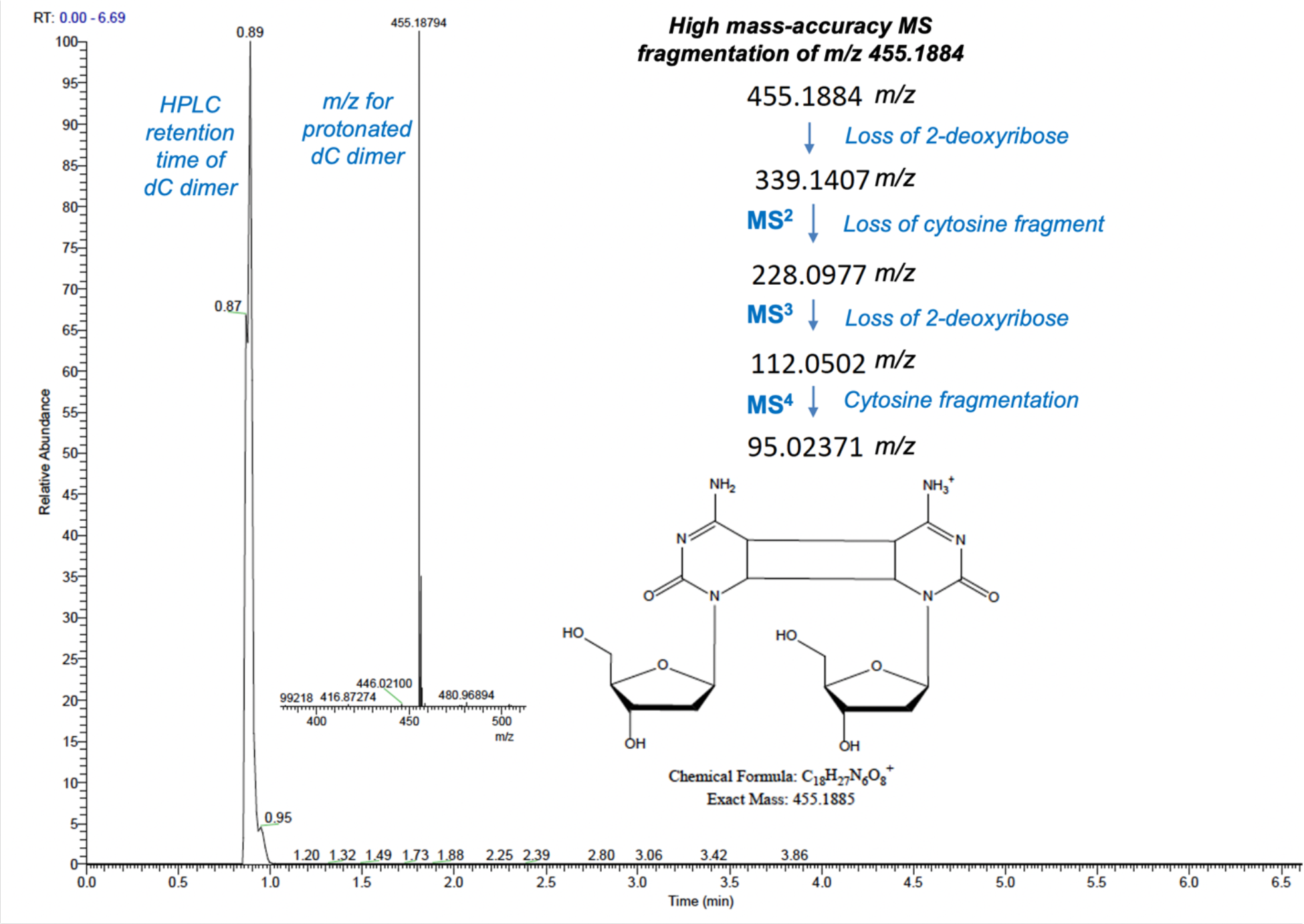
High–resolution MS^4^ Orbitrap analysis reveals the structure of the putative DNA adduct at *m/z* 455. *Left to right*: The HPLC chromatogram shows retention time of 0.89 min. The mass spectrum shows an *m/z* value of 455.18794 for the protonated species, which equates to an exact mass of 454.1801 for the adduct. The MS^4^ fragmentation series shows progressive loss of two cytosine-sized fragments and is consistent with a cyclobutane dimer of dC. These data pertain to Figure 1 and **Supplementary Table 1**.

**Extended Data Figure 3.**
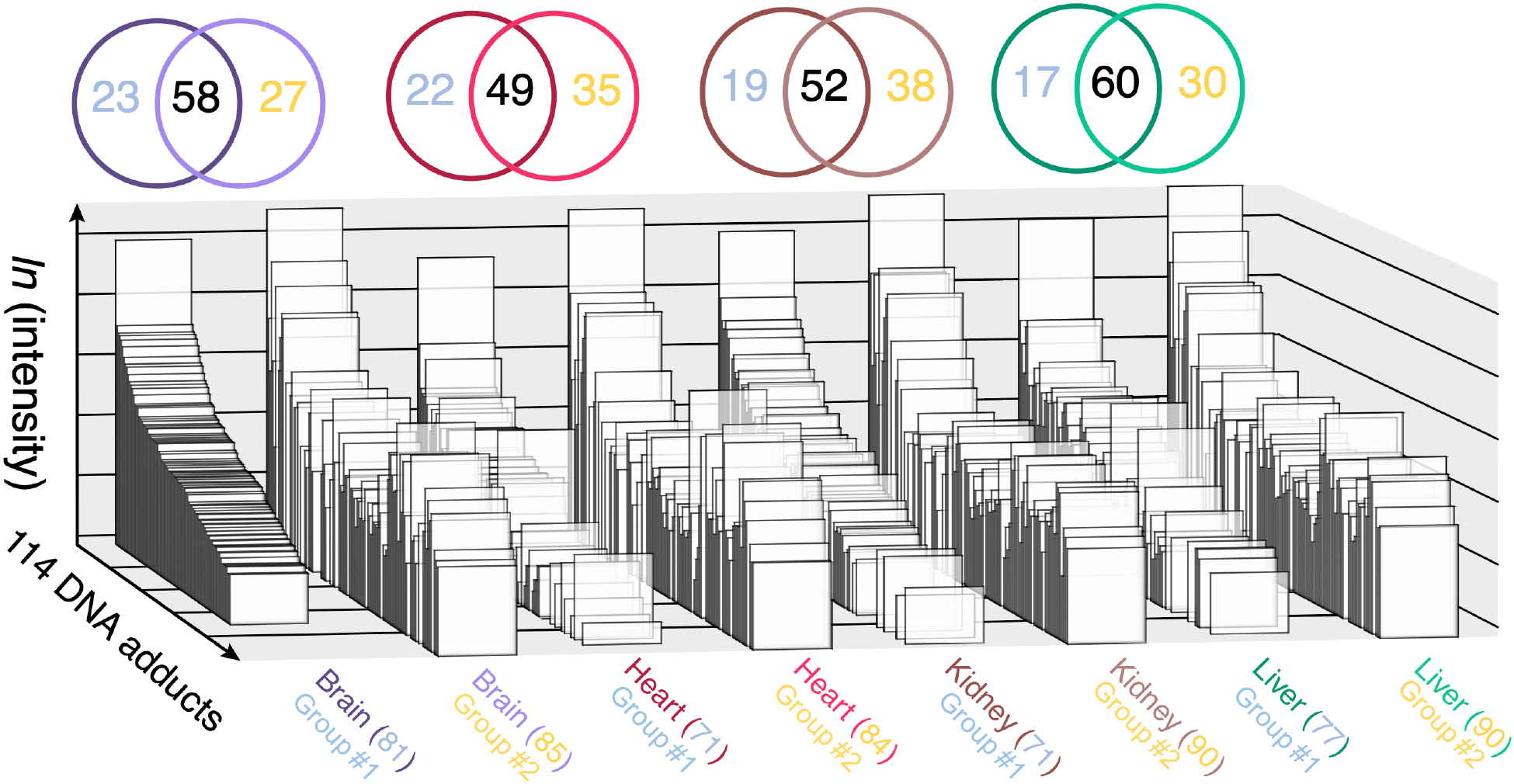
DNA damage is strongly tissue-specific in rats. Comparison of normalized LC-MS signal intensities for 92 putative DNA adducts in 4 tissues in group #1 rats (blue numbers in upper circles and lower labels) and 94 in group #2 rats (yellow numbers). Rank order was established relative to brain from group #1 (left-most plot) with comparisons to heart, kidney, and liver for group #1 and group #2 rats. For each tissue, numbers in overlapping circles correspond to unique adducts in each group of rats (blue for group #1, yellow for group #2) and shared adducts (black numbers), with matching colors of upper circles and lower labels. These data pertain to Figures 1 and 3.

**Extended Data Figure 4.**
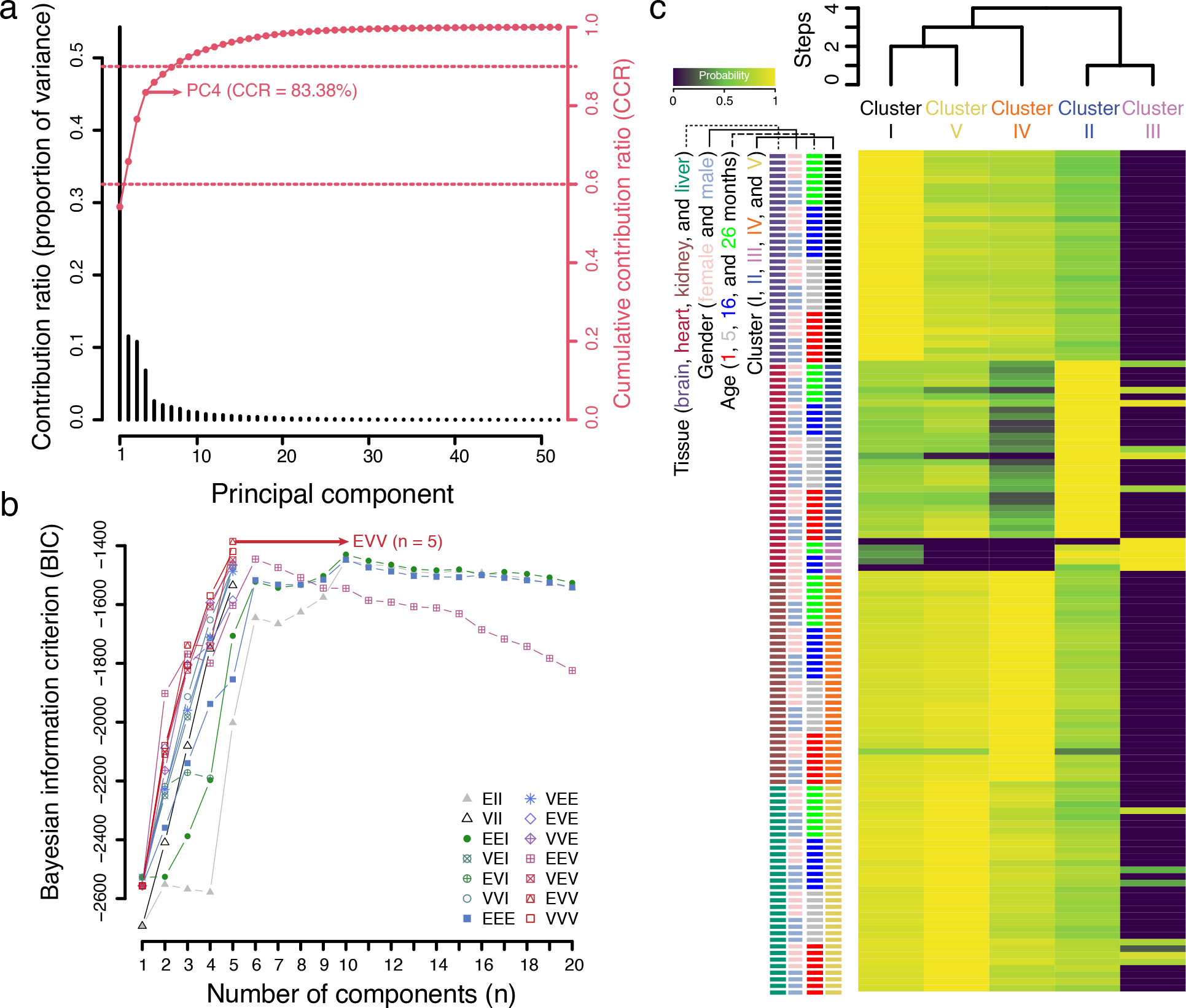
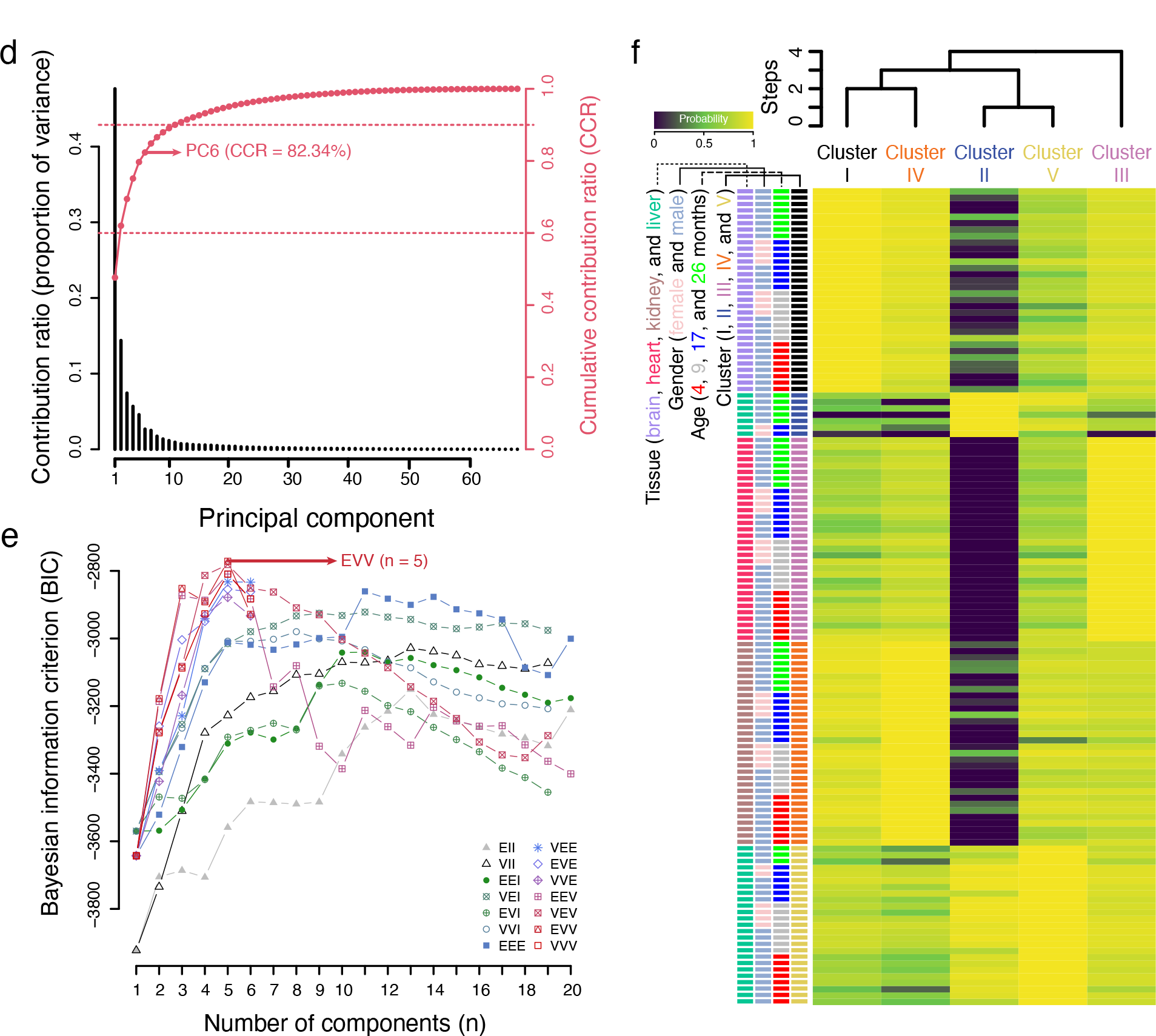
Multivariate statistical analysis of 52 (**a-c**) and 64 (**d-f**) shared DNA adducts in group #1 and group #2 rat tissues reveals 5 co-varying clusters based on age, sex, and tissue. (**a, d**) Principal components analysis identifies 2 components that account for 63% of the variance. Black bars (left axis) indicating the contribution ratio (proportion of variance), red circles (right axis) indicating the cumulative proportion of variance (cumulative contribution ratio [CCR]), and horizontal dashed lines (right axis) indicating CCRs of 60% and 90%. (b, e) Bayesian analysis to define the number of components for Gaussian mixture model (GMM) clustering. The number of components, describing the underlying Gaussian distributions, was defined based on Bayesian information criterion values of models with differing parametrizations. The best fitting model (Ellipsoidal distributions with equal volume and variable shape and orientation axes, EVV) had 5 components that were used for GMM clustering. (**c, f**) Membership probability matrix based on GMM clustering. Each value shows the posterior probability based on the model in panels **b** and **e**. The posterior probability describes the likelihood of each sample belonging to each cluster. Logarithmic transformation of conditional probabilities from expectation maximization was used to generate the heatmap. The dendrogram illustrates model-based hierarchical agglomerative clustering based on the Gaussian probability model for maximizing the resulting likelihood. This figure is associated with Figure 3b**, h.**

**Extended Data Figure 5.**
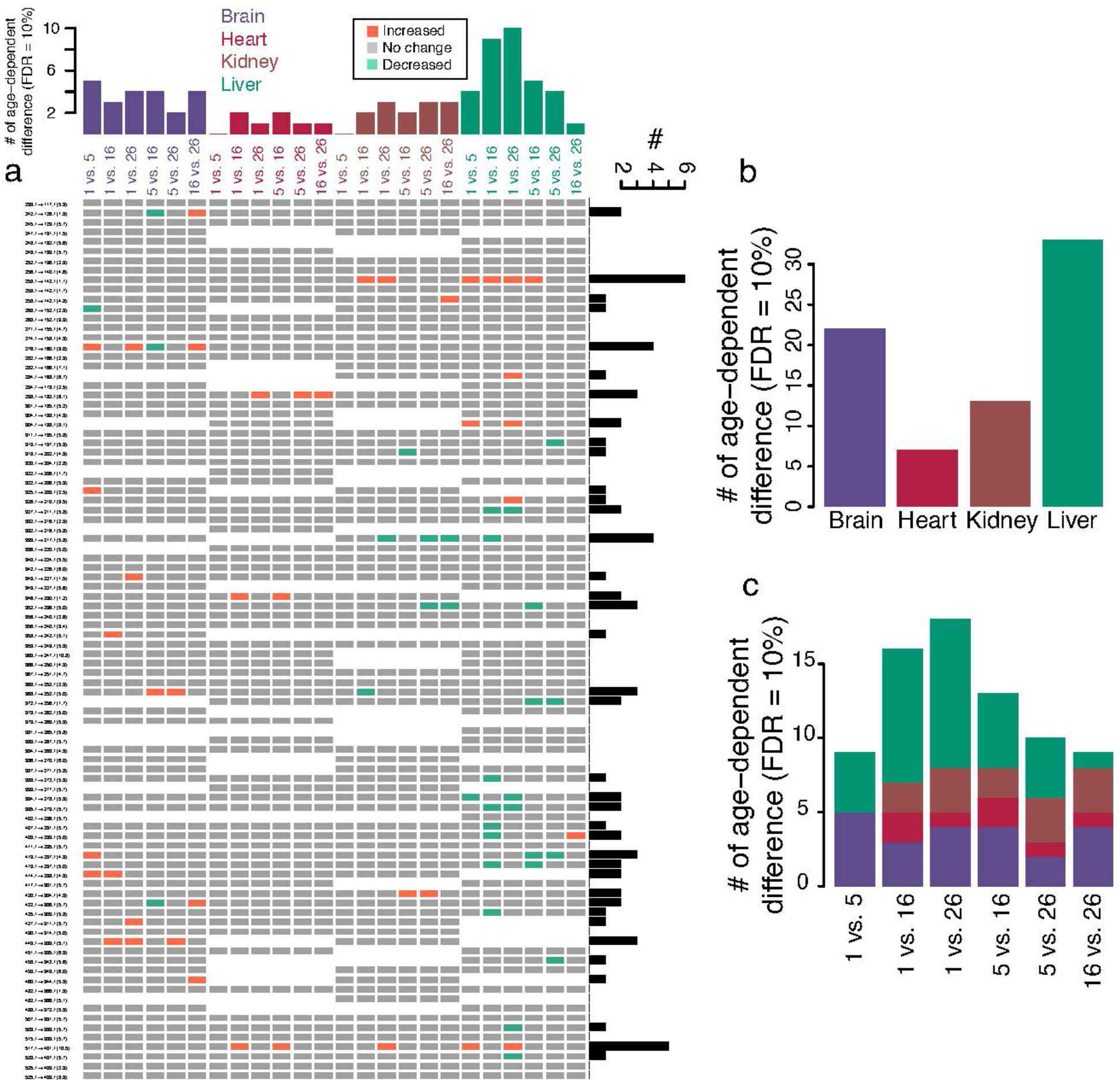
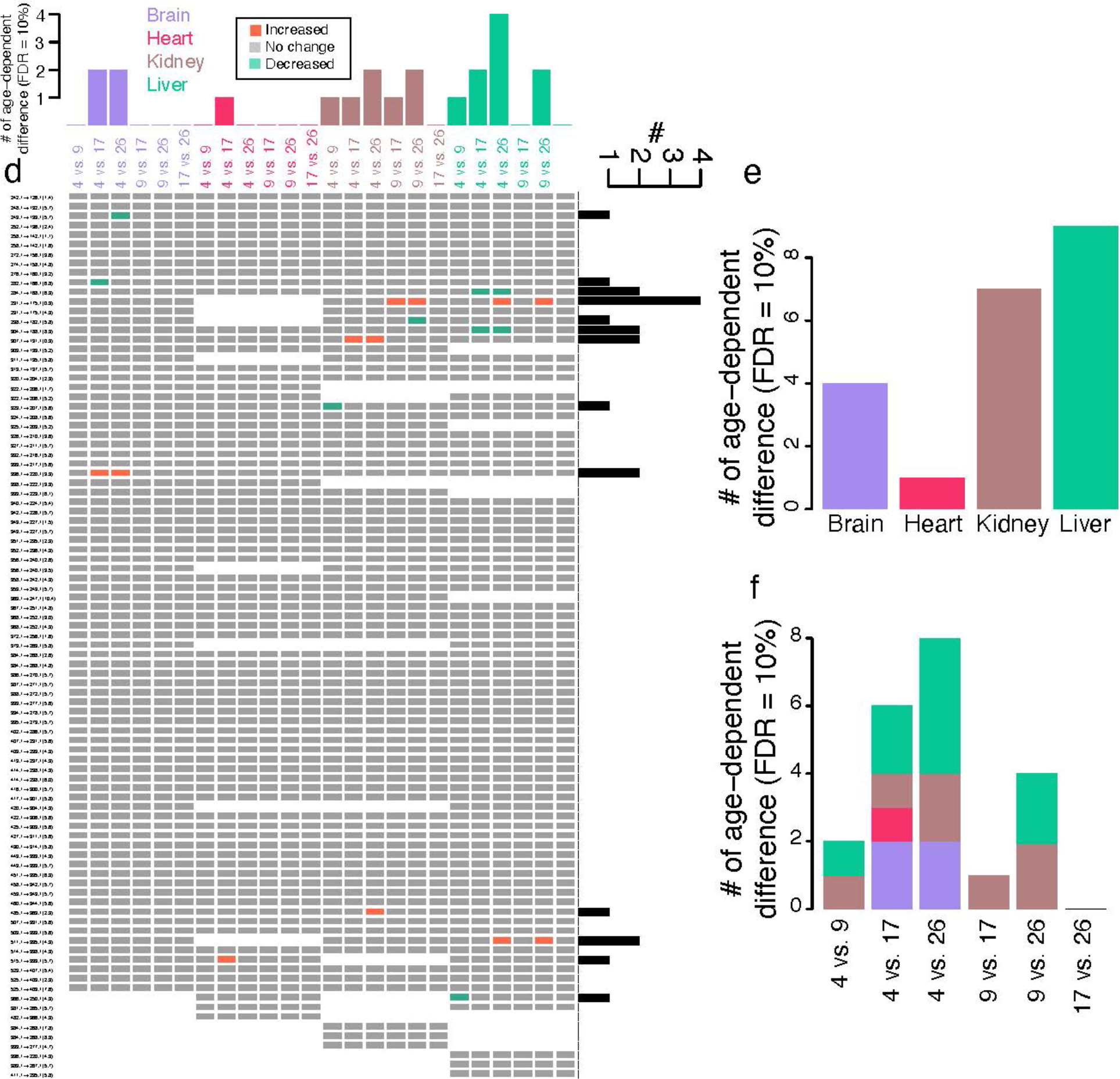
Age-biased DNA adducts in four rat tissues in two different sets of rats. **(a, d)** Heatmaps show pairwise comparisons of all ages in each tissue. Significant changes (FDR = 10%) are presented as red (increase) and green (decrease). Bar plot on the top shows the frequency in each compared age. Bar plot (black) on the right shows frequency in each adduct. (**b**, **e**) Bar plots of overall frequency of significant changes (FDR = 10%) in each tissue. (**c**, **f**) Bar plots of overall frequency of significant changes (FDR = 10%) in each age-pairwise comparison. These data pertain to Figures 1 and 4, and **Supplementary Table 5.**

**Extended Data Figure 6.**
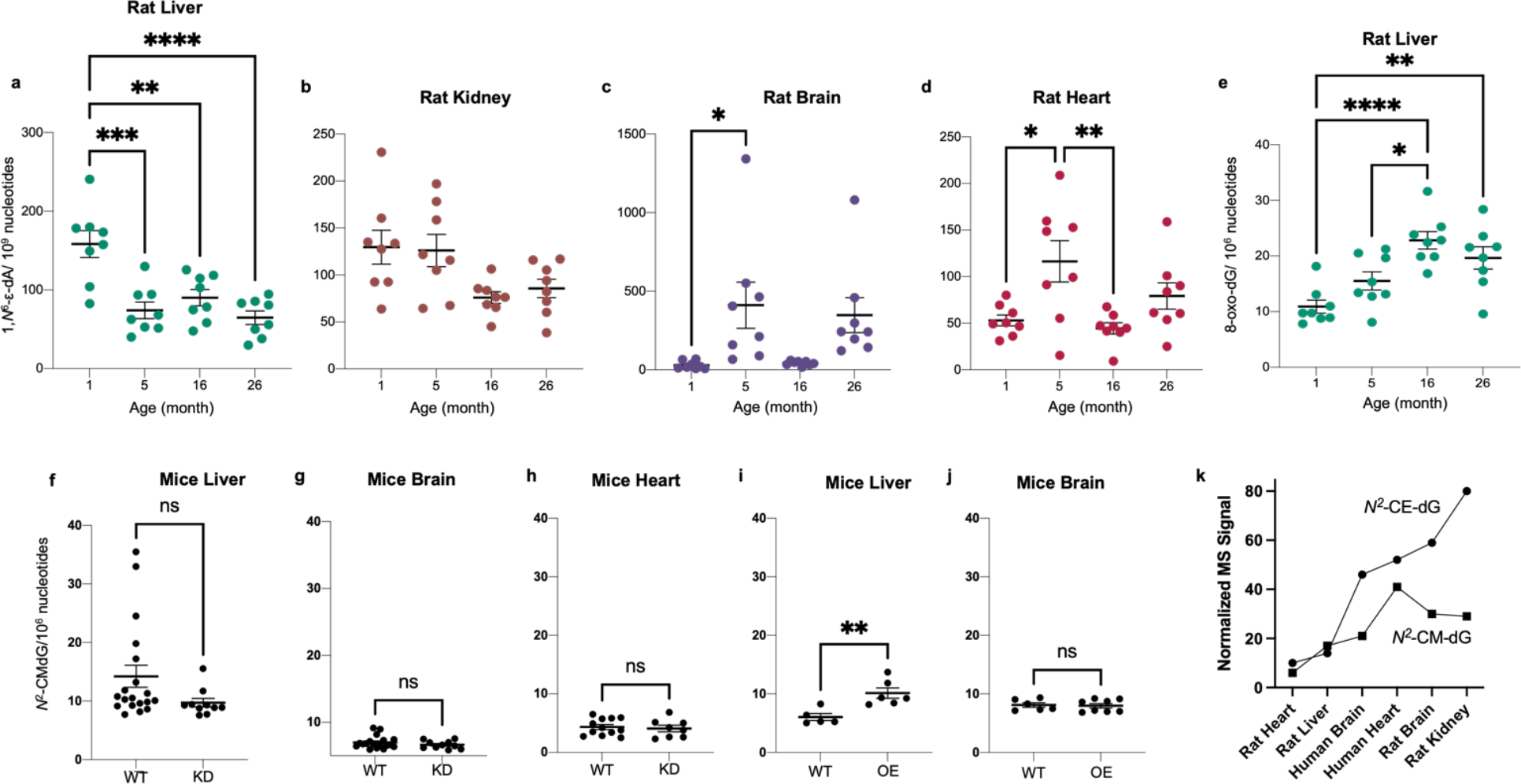
Analysis of DNA damage products in rat, mouse, and human tissues. (**a-j**) Isotope dilution chromatography-coupled triple quadrupole mass spectrometric analysis of DNA adducts in rat and mouse tissues. 1,*N*^6^-εdA levels in rat tissues as a function of age: liver (**a**), kidney (**b**), brain (**c**), and heart (**d**). Data are mean ± SEM for *n* = 8 rats. One-way ANOVA with Bonferroni’s multiple comparisons test was used to evaluate difference among ages., \**p*<0.05, \*\**p*<0.01, \*\*\**p*<0.001, \*\*\*\**p*<0.0001. (**e**) 8-Oxo-dG levels as a function of age in rat liver. Box and whisker plots show mean (heavy bar) and SEM (whiskers). One-way ANOVA with Bonferroni’s multiple comparisons test was used to evaluate differences among ages, \**p*<0.05, \*\**p*<0.01 and **** *p*<0.0001. Associated with Figure 3. (**f-j**) Glyoxalase I levels do not affect the level of *N*^2^-CMdG in transgenic mice. *N*^2^- CMdG levels were measured in liver (**f, i**), brain (**g, j**), and heart (**h**) from glyoxalase I knockdown (KD) mice (**f-h**) and from glyoxalase I over-expressing (OE) mice (**i, j**). Box and whisker plots show mean (heavy bar) and SEM (whiskers). A two-tailed unpaired Student’s *t*- test was used to evaluate the difference between wild-type (WT) and transgenic mice (KD, OE). ** *p*< 0.01, *ns*: not-significant (**f-h**). Plot of normalized MS signal intensities for putative *N^2^*-CE-dG (*m/z* 340-224) and validated *N^2^*-CMdG for rat and human tissues. Data represent mean for *n*=8 measurements of male and female tissues. This figure is associated with Figure 3.

## Extended Data Tables

**Extended Data Table 1.** Characteristics of the human tissue donors. ***Upper: heart tissue.*** LV: Left Ventricle, HMI: Heart Mass Index, LVMI: Left Ventricular Mass Index, BSA: Body Surface Area, BMI: Body Mass Index, LVEDd: Left Ventricle End-Diastolic dimension, LVEDs: Left Ventricle End-Systolic dimension, PW: Posterior Wall, LVEF: Left Ventricle Ejection Fraction. ***Lower: brain tissue*.** Donors are represented according to their ID number (#), category (control, CT), age at death and gender. *n*= 10 CT. These data pertain to Figure 5.

| Patient ID # | Age (year) | Gender | LV Mass | Heart Weight | % of LV/ Ht weight | HMI | LVMI | Weight (Kg) | Height (cm) | BSA | BMI | LVEDd | LVEDs | PW Thick | LVEF | Creatinine |
| --- | --- | --- | --- | --- | --- | --- | --- | --- | --- | --- | --- | --- | --- | --- | --- | --- |
| 1584 | 22 | Male | 182 | 299 | 61 | 164 | 100 | 67 | 180 | 1.82 | 20 |  |  |  | 55 | 1.0 |
| 1400 | 22 | Female | 168 | 278 | 60 | 145 | 88 | 88 | 150 | 1.91 | 39 | 4.1 | 2.2 |  | 65 | 0.8 |
| 1761 | 26 | Female | 170 | 254 | 67 | 137 | 92 | 69 | 180 | 1.86 | 21 | 4.0 | 3.2 | 0.9 | 50 | 2.8 |
| 1664 | 27 | Female | 165 | 260 | 63 | 137 | 87 | 79 | 165 | 1.90 | 29 |  |  |  | 58 | 1.9 |
| 1718 | 33 | Male | 189 | 278 | 68 | 157 | 107 | 65 | 173 | 1.77 | 22 | 4.9 | 3.7 | 0.7 | 55 | 0.7 |
| 1727 | 40 | Male | 198 | 300 | 66 | 145 | 95 | 90 | 172 | 2.07 | 30 | 4.9 | 3.0 | 1.3 | 60 | 3.7 |
| 1801 | 42 | Male | 208 | 377 | 55 | 206 | 114 | 69 | 175 | 1.83 | 23 | 4.4 | 2.6 | 1.2 | 62 | 0.9 |
| 1549 | 49 | Male | 213 | 383 | 56 | 168 | 93 | 104 | 180 | 2.28 | 32 |  |  |  | 55 | 2.1 |
| 1750 | 51 | Female | 150 | 227 | 66 | 148 | 98 | 51 | 167 | 1.54 | 18 | 3.8 | 3.0 | 0.9 | 60 | 1.0 |
| 1600 | 51 | Female | 134 | 213 | 63 | 122 | 76 | 68 | 163 | 1.75 | 26 | 4.2 | 2.8 | 0.7 | 50 | 0.8 |
| 1739 | 52 | Male | 213 | 352 | 61 | 168 | 102 | 90 | 175 | 2.09 | 29 | 4.1 | 2.6 | 0.8 | 65 | 0.7 |
| 1666 | 54 | Male | 159 | 262 | 61 | 152 | 92 | 62 | 173 | 1.73 | 21 |  |  | 1.2 | 65 | 0.8 |
| 1622 | 56 | Male | 156 | 259 | 60 | 154 | 93 | 66 | 155 | 1.69 | 28 | 4.2 | 2.4 | 0.9 | 65 | 1.3 |
| 1690 | 63 | Female | 137 | 247 | 55 | 153 | 85 | 56 | 167 | 1.61 | 20 | 3.6 | 2.6 | 0.8 | 60 | 0.9 |
| 1716 | 64 | Female | 158 | 269 | 59 | 139 | 82 | 81 | 166 | 1.93 | 29 | 3.9 | 2.3 | 0.9 | 70 | 0.9 |
| 1490 | 71 | Female | 104 | 206 | 50 | 142 | 72 | 48 | 158 | 1.45 | 19 |  |  |  |  | 0.5 |
| 1732 | 72 | Female | 156 | 271 | 58 | 159 | 91 | 67 | 157 | 1.71 | 27 | 4.0 | 2.9 | 0.8 | 65 | 0.6 |
| 1485 | 78 | Male | 234 | 448 | 52 | 212 | 111 | 92 | 175 | 2.12 | 30 |  |  |  |  | 3.4 |
| 1488 | 81 | Male | 218 | 402 | 54 | 210 | 114 | 78 | 170 | 1.91 | 27 |  |  |  |  | 1.6 |

| Patient ID # | Category | Age (year) | Gender |
| --- | --- | --- | --- |
| 20998065 | CT | 81.6 | Female |
| 10315029 | CT | 87.7 | Male |
| 20875195 | CT | 74.1 | Female |
| 21000504 | CT | 85.9 | Female |
| 21001933 | CT | 80.8 | Female |
| 20240514 | CT | 89.7 | Female |
| 11331231 | CT | 81.1 | Male |
| 11615242 | CT | 71.6 | Male |
| 10589255 | CT | 74.3 | Male |
| 15196262 | CT | 81.1 | Male |

## Notes

https://chorusproject.org/pages/dashboard.html#/projects/all/1767/experiments

### Competing Interest Statement

A. Palmer has a patent for the glyoxalase mice methods and use of Glo1 inhibitors (https://patents.google.com/patent/US11235020B2/en).

### Summary of Updates

Issue with figures

